# Astrocytic FKBP5 Regulates Neuroinflammation and Cognitive Outcomes in Excitotoxic Brain Injury

**DOI:** 10.64898/2025.12.02.691073

**Authors:** Yu-Ling Gan, Shang-Hsuan Lin, Yu-Ping Kang, Jia-Zhen Zhou, Wei-Hsuan Huang, Ping-Hua Sung, Chia-Chi Hung, Pei-Chien Hsu, Shu-Yin Chu, Feng-Shiun Shie, I-Hui Lee, Chung-Jiuan Jeng, Yi-Hsuan Lee

**Author notes:** Correspondence to: Yi-Hsuan Lee, Department and Institute of Physiology, National Yang Ming Chiao Tung University, No. 155, Sec. 2, Linong St. Beitou Dist., Taipei City 112304, Taiwan, Correspondence may also be addressed to: Chung-Jiuan Jeng, Department and Institute of Anatomy and Cell Biology, College of Medicine, National Yang Ming Chiao Tung University, No. 155, Sec. 2, Linong St. Beitou Dist., Taipei City 112304, Taiwan, Correspondence may also be addressed to: I-Hui Lee, Department of Neurology, Neurological Institute, Taipei Veterans General Hospital, No.201, Sec. 2, Shipai Rd., Beitou District, Taipei City 112201, Taiwan. These authors contributed equally to this work (Shang-Hsuan Lin, Yu-Ping Kang, Jia-Zhen Zhou).

## Abstract

FK506-binding protein 51 (FKBP51, encoded by *FKBP5*) is a multisignaling cochaperone that regulates cellular responses to stress. FKBP51 is upregulated in reactive astrocytes; however, the role of FKBP51 in excitotoxic brain injury remains unknown. Here, we investigated how both global and astrocyte-specific *Fkbp5* deletion influence seizure susceptibility, astrogliosis, neuroinflammation, and cognition in male mice subjected to a kainic acid (KA)-induced epilepsy mouse model. Global *Fkbp5* knockout (*Fkbp5*-KO) presented lower seizure activity along with decreased neuronal loss and astrogliosis in the hippocampus compared with the wild-type mice. Astrocyte-specific *Fkbp5* conditional knockout (a*Fkbp5*-cKO) mice similarly attenuated seizure severity, decreased astrogliosis, improved novel object recognition, and preserved glutamate transporter 1 (GLT-1) expression in hippocampal CA3. Glia-neuron mixed cultures derived from *Fkbp5*-KO brains showed reduction of NMDA-induced neurotoxicity, astrogliosis, accompanied by decreased NF-κB p65 phosphorylation. Notably, overexpression of an *Fkbp5* quadruple mutant that disrupts the FKBP51–NF-κB interaction inhibited proinflammatory lipopolysaccharide (LPS)-induced astrogliosis and NF-κB activation. The hippocampal transcriptome of the LPS-treated *Fkbp5*-KO mice revealed suppression of NF-κB signaling. In summary, this study highlights FKBP51 as a key mediator of excitotoxin-induced neuroinflammation and GLT-1 dysfunction and underlines NF-κB-mediated inflammatory astrogliosis as a potential intervention target for excitotoxic brain injury.

## 1. Introduction

FK506 binding protein 51 (FKBP51, 51 kDa), encoded by the *FKBP5* gene, belongs to the tetratricopeptide repeat protein-containing immunophilin family (Schiene-Fischer and Yu, 2001; Wiederrecht et al., 1992). FKBP51 is best known as a heat shock protein 90 (Hsp90)-associated cochaperone that regulates the activity of the glucocorticoid receptor (GR) (Pratt, 1993; Wochnik et al., 2005). FKBP51-bound GR exhibits low cortisol affinity and increased resistance to steroid hormones and exhibits reduced nuclear transactivation of glucocorticoid response element (GRE)-containing genes, including *FKBP5*(de Kloet et al., 2005; Muzikar et al., 2009). Upon stress and hypothalamic pituitary adrenal (HPA) axis activation (Binder, 2009; Zannas et al., 2016), cortisol levels increase, and FKBP51 is displaced by other chaperones, leading to GR nuclear transactivation. *FKBP5* gene expression is then upregulated via GRE, creating a negative feedback loop that decreases the sensitivity of GR to cortisol. The intronic rs1360780 risk allele T, a well-studied single nucleotide polymorphism (SNPs) of the *FKBP5* gene, promotes allele-specific DNA demethylation and stress-driven *FKBP5* overexpression (Klengel et al., 2013), increasing vulnerability to neuropsychiatric disorders, including posttraumatic stress disorder and major depression (Han et al., 2017; Storer et al., 2011; Yehuda et al., 2013).

FKBP51 is a multifaceted regulator of biological processes and signaling pathways, including metabolism (Balsevich et al., 2017), neuronal signaling (Hausch, 2015), and nuclear factor kappa B (NF-κB) signaling (Erlejman et al., 2014; Romano et al., 2015). NF-κB is a transcription factor that governs the expression of numerous proinflammatory genes encoding cytokines, chemokines, and adhesion molecules (Liu et al., 2017a). In the resting state, NF-κB is held inactive in the cytoplasm by IκB. Upon activation during inflammation, the IκB kinase (IKK) complex phosphorylates IκB, which targets it for degradation. This process releases NF-κB, allowing it to translocate to the nucleus and activate the transcription of proinflammatory genes. In the central nervous system (CNS), excessive NF-κB activation in microglia and astrocytes often exacerbates neurotoxicity (Kwon and Koh, 2020). Our previous studies demonstrated that FKBP51 contributes to neuroinflammation by regulating NF-κB signaling in both in vitro primary microglial cultures and in vivo *Fkbp5* knockout (*Fkbp5*-KO) mice. Following lipopolysaccharide (LPS)-induced CNS and peripheral inflammation, *Fkbp5* deletion attenuates microglial activation and decreases NF-κB activation, accompanied by a reduction in the proinflammatory cytokine TNF-α and inflammatory mediators such as inducible nitric oxide synthase and cyclooxygenase-2. These findings highlight the critical role of FKBP51 in regulating CNS inflammation (Gan et al., 2024; Gan et al., 2022). Interestingly, LPS increases glutamic acid decarboxylase 65 (GAD65), a membrane-associated GABA-synthesizing enzyme, in the hippocampus of wild-type (WT) but not *Fkbp5*-KO mice. This FKBP51-dependent GAD65/GABA increase is observed in the ventral hippocampus and is associated with resilience to inflammation-induced anxiety-like behaviors. These findings suggest that FKBP51 is a crucial regulator of neuroinflammation and neuroexcitability (Gan et al., 2022).

Neuroexcitotoxicity serves as a central mechanism driving neuronal damage in epilepsy. Given that stress-induced HPA axis activation exacerbates seizures in epilepsy (Maguire and Salpekar, 2013), we hypothesized that stress-induced upregulated FKBP51 expression contributes partly to brain excitotoxic injury. Prolonged excitotoxicity triggers the pathological overactivation of glutamate receptors and excessive release of excitatory neurotransmitters (Haroon et al., 2017; Sofroniew, 2014) and inflammatory cytokines, resulting in a self-propagating cycle of inflammation and neurotoxicity (Dong et al., 2009; Hatton et al., 2020; Wang et al., 2016). Astrocytes play a critical role in maintaining CNS homeostasis by regulating ion balance, clearing excess glutamate, and supporting neurotransmission. However, astrocytic dysfunction, including impaired glutamate transporter-1 (GLT-1) activity, disrupts glutamate clearance, leading to excitotoxic damage and increased susceptibility to seizures (Ben Haim and Rowitch, 2017; Haroon et al., 2017; Pajarillo et al., 2019; Rana and Musto, 2018). We previously showed that the preservation of GLT-1-expressing perineuronal astrocytes in the hippocampal CA3 subregion reduces excitotoxic kainic acid (KA)-induced seizures, memory impairment, and neuronal loss (Kuo et al., 2019). Recent transcriptomic studies have identified FKBP51 as an enriched marker in reactive astrocytes (Clarke et al., 2018; Liddelow et al., 2017). However, whether FKBP51 mediates astrocytic responses in the context of brain excitotoxic injury is unknown.

Based on our findings that FKBP51 regulates the NF-κB signaling pathway in neuroinflammation (Gan et al., 2024; Gan et al., 2022) and that NF-κB is a driver of proinflammatory responses in astrocytes (Guo et al., 2024; Linnerbauer et al., 2020), we hypothesized that astrocytic FKBP51 deficiency offers neuroprotection in excitotoxic brain injury. To test this hypothesis, we employed total *Fkbp5*-KO and conditional *Fkbp5* deletion in astrocytes in a KA-induced epilepsy mouse model (Vincent and Mulle, 2009). We assessed seizure severity, neuronal loss, and astrogliosis in vivo, and performed mechanistic studies in primary glia–neuron mixed cultures to elucidate the role of FKBP51 in reactive astrogliosis. This study highlights FKBP51 is a potential therapeutic target for epilepsy and other excitotoxic disorders.

## 2. Materials and methods

### 2.1. Animals

Male C57BL/6J (wild-type, WT), *Fkbp5^tm1Dvds^*/J (*Fkbp5*-KO; Jackson Laboratory, Stock No. 017989), *Fkbp5*-floxed (*Fkbp5*^fl/fl^), and *Slc1a3-*Cre^ERT^;*Fkbp5*^fl/fl^ mice were used. *Fkbp5*-KO mice and *Slc1a3*-Cre^ERT^ transgenic mice (Tg (*Slc1a3*-cre/ERT)1Nat/J; Stock No. 012586) were obtained from Jackson Laboratory. *Fkbp5*^fl/fl^ mice were generated using CRISPR/Cas9 to insert two loxP sequences into exon 3 of the *Fkbp5* gene, performed by the Transgene Core Facility at the Institute of Molecular Biology (IMB, Academia Sinica, Taiwan). To generate inducible astrocyte-specific conditional *Fkbp5*-KO (a*Fkbp5*-cKO) mice, *Fkbp5*^fl/fl^ mice were mated with *Slc1a3*-Cre^ERT^ mice to generate *Slc1a3*-Cre^ERT^;*Fkbp5*^fl/+^ offspring, which were then bred with *Fkbp5*^fl/fl^ mice to generate *Slc1a3-*Cre^ERT^;*Fkbp5*^fl/fl^ mice. Each individual mouse was considered an experimental unit. Genotyping of the floxed *Fkbp5* and *Slc1a3*-Cre^ERT^ alleles was performed using PCR. Primer sequences are listed in Supplementary Table S1. The integrity of loxP and Cre transgene was verified by agarose gel electrophoresis. At 5 weeks of age, male *Fkbp5*^fl/fl^ and *Slc1a3-*Cre^ERT^;*Fkbp5*^fl/fl^ mice received intraperitoneal tamoxifen injections (100 mg/kg/day for 5 consecutive days; Sigma Aldrich) to induce ERT-driven Cre-mediated *Fkbp5* deletion in GLAST-expressing astrocytes (Huang et al., 2023). At 12 weeks of age, mice were randomly assigned to experimental groups, male mice received intracerebroventricular or intraperitoneal injections of kainic acid (KA) or saline. The study included four groups: WT, *Fkbp5*-KO, *Fkbp5*^fl/fl^, and a*Fkbp5*-cKO, with saline-injected mice serving as controls. The induction of astrocyte-specific *Fkbp5* deletion in a*Fkbp5*-cKO mice was validated by isolating astrocytes. The astrocyte isolation procedure was modified from a previous report (Huang et al., 2023) and used an anti-ACSA-2 antibody (Miltenyi Biotec) conjugated with magnetic beads for separation, followed by Western blot for FKBP51 and GLAST expression in the sorted cells. Animals were excluded if they lacked the required genotype or died from kainic acid (KA)-induced seizures before the endpoint. Mice bred in the Laboratory Animal Center of National Yang Ming Chiao Tung University, Taipei, Taiwan. The mice were fed a chow diet and kept in a 12-h light/dark cycle at 25L±L2 °C. All animal procedures were approved by the Institutional Animal Care and Use Committee (IACUC) of National Yang Ming Chiao Tung University (IACUC No.: 1090515 and 1100336). Sample sizes were chosen based on our previous studies employing similar tests (Gan et al., 2022; Kuo et al., 2019).

### 2.2. Intracerebroventricular and intraperitoneal injection of KA

Kainic acid (KA) is a potent neurotoxin that can induce neuronal damage and epileptic seizures resembling temporal lobe epilepsy (Vincent and Mulle, 2009; Zheng et al., 2011). The intracerebroventricular (icv) injection of KA was performed in 12-week-old male C57BL/6J or *Fkbp5*-KO mice to induce excitotoxic injury for 24 hours. In brief, the mice were anesthetized by the intraperitoneal injection of avertin (100 mg/kg body weight). The anesthetized mice were placed in a stereotaxic device (David Kopf Instruments), and an injection needle was surgically inserted at the following coordinates with reference to bregma: 0.4 mm posterior to bregma, 1 mm lateral to the midline, and 2.2 mm under the dura. Sterile saline or KA (Millipore, Cat. No. 420318; 0.5 μg/2 μl of saline) was injected into the left and right lateral ventricles at a constant infusion rate of 1 μl/min over a 2-minute period using an infusion pump (KD Scientific). For the intraperitoneal injection of KA, 12-week-old male mice were administered 20 mg/kg body weight KA for 7 days. The mice were sacrificed and intracardially perfused with saline. Brains were collected for Western blot, RT qPCR, and immunofluorescence.

### 2.3. Assessment of seizure activity

Seizure activity, assessed as the primary outcome using a modified Racine scale (Benkovic et al., 2004; Kuo et al., 2019), was the scored by blinded observers: (1) freezing behavior; (2) rigid posture with raised tail; (3) continuous head bobbing and forepaw shaking; (4) rearing, falling, and jumping; (5) continuous level 4 activity; and (6) loss of posture and generalized convulsions. Seizures were monitored from 1.5–5.5 hours and at 24 hours after intracerebroventricular injection of KA, or from 10–120 minutes after intraperitoneal injection of KA.

### 2.4. Immunofluorescence staining and confocal microscopy

Mice were transcardially perfused with saline, and brains were postfixed in 4% paraformaldehyde for 24 hours, then cryoprotected in 30% (w/v) sucrose. Coronal brain sections (30-μm thickness) were permeabilized, incubated with primary and secondary antibodies conjugated with Alexa Fluor (Invitrogen), and coverslipped using fluorescent mounting medium with DAPI (Vector Laboratories). For cultured cells, coverslips were fixed and processed as described above. Antibodies used are listed in Supplementary Table S2. Images were captured using a Leica DM 6000B fluorescence microscope or an Olympus FV1000 laser confocal microscope. Signal intensity and expression areas were quantified using MetaMorph version 7.8 software (Molecular Devices). The number of NeuN-positive cells was counted and normalized to the total number of DAPI-stained cells. The GFAP and GLT-1 areas were quantified on the basis of their immunoreactive area and normalized to the respective region of interest. Nodes of Ranvier were defined by the presence of two Caspr proteins flanking a Na_V_1.6-positive site, representing a single node (Caldwell et al., 2000). Node density was calculated as number per 1 mm², and Na_V_1.6 domain length was measured as distance between the Caspr-labeled regions. The percentage of MAP2-circled neurons was calculated by determining the ratio of MAP2-positive cells to the total number of DAPI-stained cells.

### 2.5. Western blot analysis

Brain tissues or cells were lysed as previously described (Hung et al., 2016). Protein samples were resolved by SDS-PAGE, transferred to membranes, and incubated with primary and HRP-conjugated secondary antibodies. Signals were detected using enhanced chemiluminescence reagent (Western Lightning Plus-ECL, PerkinElmer) and visualised with ImageQuant LAS 4000 (GE Healthcare Bio-Sciences AB). Band intensity was quantified using ImageJ software (National Institutes of Health, Bethesda) and normalized to GAPDH. Antibodies used are listed in Supplementary Table S2. Uncropped images of all Western blots are provided in Supplementary Fig. S1 and S2.

### 2.6. Animal behavioral assessment

The open field test (OFT) was used to assess locomotor activity and anxiety-like behavior. Mice were placed in a 26 × 26 × 30 cm arena, and movement was tracked for 5 minutes using Smart 3.0.0.6 software (Panlab). Total distance traveled indicated locomotor activity, while the percentage of time in the central zone reflected anxiolytic behavior, indicating reduced anxiety. The novel object recognition (NOR) test was used to assess memory performance based on the natural preference of rodents for novel objects (Goulart et al., 2010; Silvers et al., 2007). Prior to testing, the mice were placed in a at the center of a 26 × 26 × 30 cm arena and allowed to freely explored two identical objects placed in opposite corners for three consecutive days. On the fourth day, one of the objects was replaced with a novel object, and mice were allowed to explore freely for 5 minutes. Interaction time and movement trajectories toward the object were recorded. The discrimination index, a metric used to quantify memory performance, was calculated as: discrimination index = time spent exploring novel objects/(time spent exploring novel objects + time spent exploring familiar objects). All tests were conducted in a controlled behavioral room under consistent lighting during the light phase. Cage location and injection order were randomized to minimize confounding. Primary outcomes were total distance (OFT) and discrimination index (NOR). Behavioral assessments were performed by experimenters blinded to group allocation.

### 2.7. Primary culture of mouse brain cells

Primary mouse mixed glia**–**neuron (GN) cultures were prepared from postnatal day 0 (P0) mouse cerebral cortices as previously described (Kuo et al., 2019), using 50% DMEM (Gibco) supplemented with 10% FBS (Invitrogen) and 50% neurobasal medium supplemented with B27. After 9 DIV, GN cultures yielded approximately 15% neurons and 85% astrocytes, characterized by MAP2 and GFAP immunostaining (Kuo et al., 2019). To induced excitotoxicity, GN cultures were treated with NMDA (5 µM; Sigma Aldrich, Cat. No. M3262). Primary mixed glial cultures were prepared from P1–P2 WT and *Fkbp5*-KO mice following a previously described protocol (Huang et al., 2023), using DMEM/F12 (Gibco) supplemented with 10% FBS, 100 U/ml penicillin and 100 μg/ml streptomycin. After 10L14 days, astrocytes were isolated by orbital shaking at 180 rpm for 2 hours at 37 °C, followed by trypsinization and centrifugation at 300 × g for 5 minutes and reseeded for experiments. To induce inflammatory responses, astrocytes were treated with LPS (1 µg/mL; *Escherichia coli* O55:B5, Sigma Aldrich, Cat. No. L5418).

### 2.8. Molecular modeling for protein***L***protein docking

Protein protein docking of FKBP51 and IKKα was performed by Dr. Chia-Cheng Chou at the National Laboratory Animal Center (Taipei, Taiwan). The crystal structures of FKBP51 (PDB: 5NJX) and IKKα (PDB: 5EBZ) were downloaded from the Protein Data Bank (Berman et al., 2000). Three structural datasets of FKBP51/IKKα were applied to the ClusterPro server for docking analysis. The docking procedure consisted of three main steps. First, rigid body docking was performed by sampling billions of conformations to identify potential binding interactions between FKBP51 and IKKα. Second, the 1000 lowest-energy structures were clustered on the basis of root-mean-square deviation (RMSD), with the largest clusters representing the most likely models of the FKBP51LIKKα complex. Finally, the selected structures underwent energy minimization to optimize the binding interfaces and validate the conformations. This modeling approach provided predictions of the key interaction sites between FKBP51 and IKKα, which were used for further mutagenesis and functional assays.

### 2.9. Fkbp5 plasmid construction and transfection

To investigate the interactions of FKBP51 with its key binding target, IKKα, molecular modeling and binding site prediction were performed by Dr. Chia-Cheng Chou at the National Laboratory Animal Center (Taipei, Taiwan). On the basis of those predictions, we first constructed a plasmid containing the mouse *Fkbp5* wild-type sequence (5.4 kb for the coding region), which was inserted into the EcoRI and XbaI restriction sites of the pcDNA3.1-Myc vector (Myc-*Fkbp5*-WT). Site-directed mutagenesis to generate the FKBP51 quadruple mutant (Myc-*Fkbp5*-3AR; Y302A, E301A, E258A, or G345R), designed to disrupt FKBP51–IKKα binding, was conducted by the Biomedical Resource Core of the First Core Laboratory, College of Medicine, National Taiwan University. Plasmid amplification were performed using Presto™ Mini and Presto™ Midi Plasmid kits (Geneaid). Primary astrocytes were transfected with the plasmids using Lipofectamine™ 3000 (Thermo Fisher Scientific) following the manufacturer’s protocol.

### 2.10. Reverse transcription***L***quantitative PCR (RT***L***qPCR)

Total mRNA was extracted using TRIzol reagent (Invitrogen) in accordance with the manufacturers’ instructions and reverse transcribed into cDNA using a high-capacity reverse transcription kit (Thermo Fisher Scientific). Quantitative PCR was performed using an ABI StepOnePlus real-time PCR system (Applied Biosystems). Relative fold change in gene expression was determined using the ΔΔCt method, normalized to 18S rRNA, and expressed as 2^-ΔΔCt^. Primer sequences are listed in Supplementary Table S3.

### 2.11. RNA sequencing and ingenuity pathway analysis (IPA)

RNA sequencing (RNA-seq) was conducted to characterize and analyze the transcriptomes of hippocampal tissues from WT and *Fkbp5*-KO mice 7 days after LPS injection (3 mg/kg; WT-SAL, WT-LPS, *Fkbp5*-KO-SAL, and *Fkbp5*-KO-LPS). Total RNA was extracted and submitted to BIOTOOLS Co., Ltd. (Taiwan) for sequencing on the NovaSeq 6000 platform using 150 bp paired-end reads. Raw read counts were obtained and used as inputs for differentially expressed gene (DEG) analysis between the groups using the DESeq2 package. Genes with a *P* value < 0.05 and a |log2-fold change| > 0.5 were considered significantly differentially expressed. Gene set enrichment analysis (GSEA) was performed using the Hallmark and Gene Ontology (GO) Biological Processes gene sets to identify enriched pathways between the WT-LPS and WT-SAL groups and between the *Fkbp5*-KO-LPS and WT-LPS groups, with significance determined by the p value and normalized enrichment score (NES). An NES > 1 indicated an activated pathway, whereas an NES < -1 indicated a suppressed pathway. Further analysis of upstream regulators was conducted using an Ingenuity Pathway Analysis (IPA) system (QIAGEN Digital Insights), with an overlapping *p* value of *p* < 0.05 used as the threshold for significance, with Z score > 1 indicating the activation of upstream regulators and Z score < -1 indicating the inhibition of upstream regulators.

### 2.12. Statistical analysis

Statistical analysis was performed using GraphPad Prism 6 software. Data are expressed as means ± SEMs. Two-way repeated measures ANOVA with Bonferroni’s post hoc test used for seizure score analysis. Two-way ANOVA with Tukey’s multiple comparisons test was used for group comparisons of expression data. One-way ANOVA with Tukey’s post hoc test was used to compare multiple groups. Unpaired Student’s *t* test was used for two-group comparisons. Full statistical results are provided in Supplementary Table S4.

## 3. Results

### 3.1. Fkbp5 deletion attenuates excitotoxin-induced seizure activity, neuronal loss, astrogliosis, and white matter injury

To investigate the role of FKBP51 in excitotoxicity, we used intracerebroventricular (icv) injection of KA (KA-icv) at a dose of 0.5 μg/mouse in wild-type (WT) and *Fkbp5*-KO mice to assess the seizure activity following the injection and brain immunohistochemistry at 24 h (Fig. 1A). Both WT and *Fkbp5*-KO mice exhibited progressive seizure behaviors following KA injection. The seizure scores for the KA-icv-treated *Fkbp5*-KO mice were significantly lower than those of WT mice, particularly between 3.5 and 5 hours after KA injection (Fig. 1B). This resilience to excitotoxicity in *Fkbp5*-KO mice was validated by brain pathology with the neuron marker NeuN and the reactive astrocyte marker GFAP. The *Fkbp5*-KO mice had significantly preserved NeuN-positive neurons in the CA1 region but not in the CA3 region (Fig. 1C, D). Notably, KA-induced GFAP-positive astrogliosis in the CA1 and CA3 regions was evident only in WT mice, not in *Fkbp5*-KO mice (Fig. 1E, 1F). These results indicate that *Fkbp5* deletion reduces susceptibility to KA-induced seizure activity, neuronal damage, and reactive astrogliosis in the hippocampus.

**Fig. 1.**
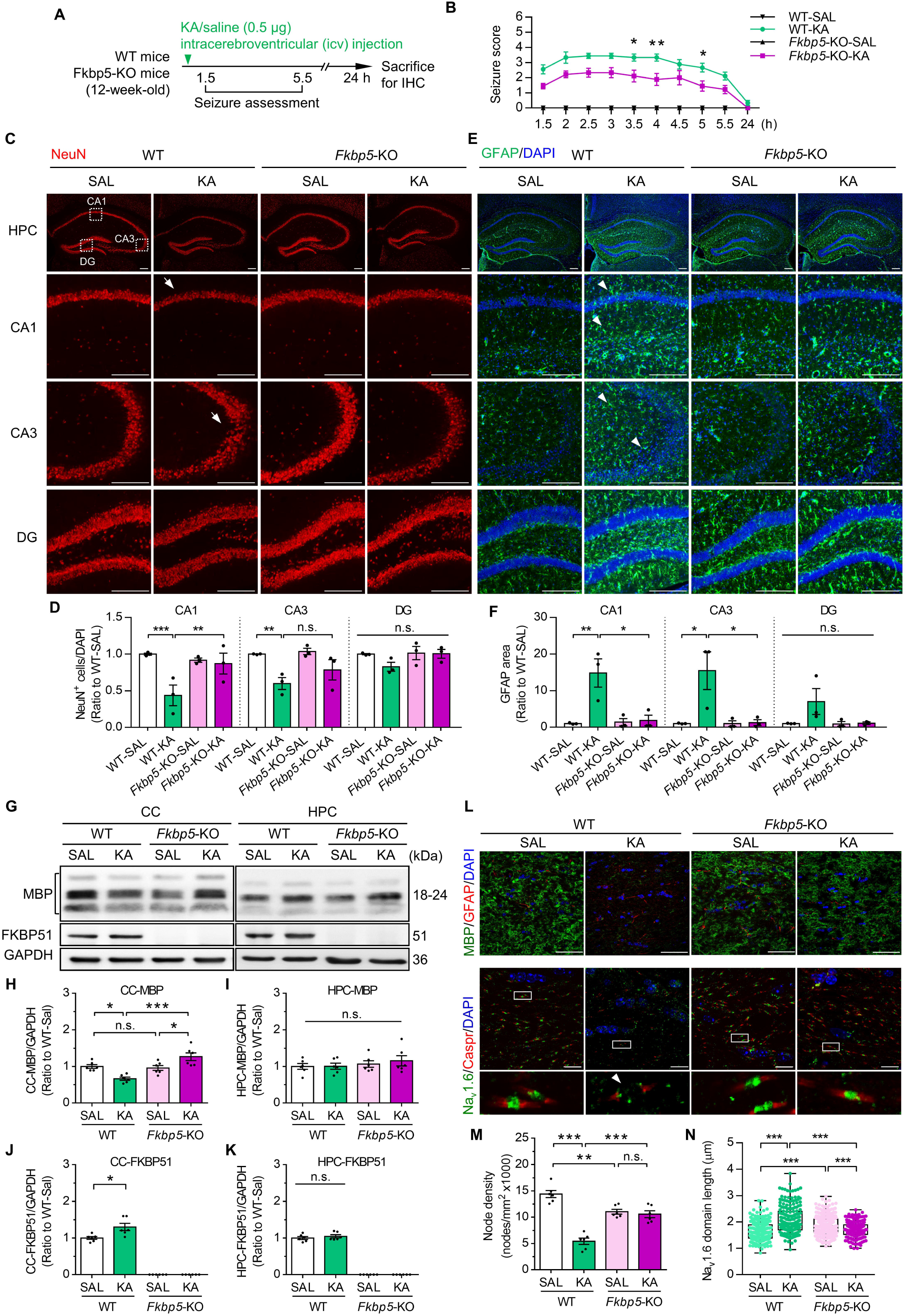
*Fkbp5*-KO mice exhibit resistant to KA-induced seizures, hippocampal damage, astrogliosis, and white matter demyelination. (**A**) Experimental timeline of intracerebroventricular injections of saline (SAL) or kainic acid (KA, 0.5 μg) in 12-week-old male wild-type (WT) and *Fkbp5*-KO mice. (**B**) Seizure score was assessed after KA injection at the indicated time points (mean ± SEM, *n* = 9 per group). * *p* < 0.05 and \*\**p* < 0.01 for the comparison between two groups in a two-way repeated measures ANOVA followed by Bonferroni’s post hoc test. (**C, E**) Representative immunofluorescence images of NeuN (green) and GFAP (red) counterstained with DAPI in the hippocampus (HPC, upper panel) and higher magnification of hippocampal CA1, CA3 and dentate gyrus (DG) subregions (lower panel). Scale bar: 100 μm. Arrow: NeuN-positive cell loss. Arrowhead: GFAP-positive increase and cellular hypertrophy. (**D, F**) Quantification of NeuN-positive cells per DAPI and GFAP area in CA1, CA3, and DG (*n* = 3 per group). (**G**) Western blot analysis of MBP and FKBP51 expression in the corpus callosum (CC) and HPC. GAPDH was used as the loading control. (**H**–**K**) Quantification of MBP and FKBP51 expression (*n* = 6 per group). (**L**) Representative immunofluorescence images of MBP (green) and GFAP (red) counterstained with DAPI in the CC (upper panel, scale bar: 50 μm) and Na_V_1.6 (green) and Caspr (red) counterstained with DAPI in the CC, with a zoomed-in view of the indicated region (lower panel, scale bar: 10 μm). Arrowhead: decreased node density and increased Na_V_1.6 domain length. (**M, N**) Quantification of node density per 1 mm^2^ (*n* = 6 per group) and Na_V_1.6 domain length (*n* = 205–278 per group). Data are expressed as the mean ± SEM. In D, F, H, and I, \**P* < 0.05, \*\**P* < 0.01, and \*\*\**P* < 0.001 for the comparisons between two groups using two-way ANOVA followed by Tukey’s post hoc test. In J, K, \**P* < 0.05 for comparisons between groups using an unpaired Student’s *t* test.

To further investigate the excitotoxicity of the white matter (WM), we examined the expression of myelin basic protein (MBP), which represents the myelin integrity of the corpus callosum and hippocampus. Western blot analysis revealed that MBP levels were significantly lower in the corpus callosum of KA-treated WT mice, but not in *Fkbp5*-KO mice (Fig. 1G, H); whereas MBP levels in the hippocampus were similar between groups (Fig. 1G, I). Notably, KA-icv administration resulted in an increase in FKBP51 protein specifically in the corpus callosum of WT mice but not in the hippocampus (Fig. 1G, J, K), suggesting that excitotoxin-induced demyelination of the brain WM might be attributed to an increase in FKBP51. Indeed, *Fkbp5*-KO mice presented preserved MBP levels (Fig. 1L). We next examined the integrity of the nodes of Ranvier via the nodal-enriched sodium channel Na_V_1.6, which was labeled together with the paranodal contactin-associated protein Caspr. Each pair of Caspr-positive segments flanking the Na_V_1.6-positive nodal segment was counted as one node (Fig. 1L). Quantitative analysis revealed a marked reduction in nodal density in the corpus callosum of KA-icv-treated WT mice (Fig. 1M), as well as a significant increase in Na_V_1.6-labeled nodal length, compared with those observed in saline-treated WT mice (Fig. 1N), indicating early demyelination characterized by axon myelin decoupling that allowed nodal proteins to diffuse into the paranodal region. Interestingly, in the *Fkbp5*-KO group, a KA-induced decrease in nodal density and increase in nodal length were not observed.

To assess the role of FKBP51 in oligodendrocyte (OL) susceptibility to excitotoxicity, we analyzed primary OL cultures exposed to AMPA, an excitotoxic mediator in OLs (Supplementary Fig. S3). In WT OLs, AMPA treatment induced *Fkbp5* expression and reduced *Mbp* and erythropoietin (*Epo*, a key OL survival factor) expression. In contrast, *Fkbp5*-KO OLs maintained *Mbp* and *Epo* expression, an effect that was associated with preserved OL morphology and reduced apoptosis. These findings suggest that FKBP51 exacerbates excitotoxic OL damage and suppresses EPO signaling, which might contribute to demyelination in vivo.

Together, these results indicate that FKBP51 exacerbates KA-induced seizures, neuronal loss and astrogliosis, while its deletion confers resilience to hippocampal and white matter injury, highlighting FKBP51 as a key driver of excitotoxicity.

### 3.2. Generation and phenotypes of astrocyte-specific Fkbp5 conditional knockout mice (aFkbp5-cKO)

The FKBP51-dependent astrogliosis observed in the hippocampus of KA-treated mice prompted us to investigate the specific role of FKBP51 in the astrocytic response to excitotoxicity. We generated astrocyte-specific *Fkbp5* conditional knockout (a*Fkbp5*-cKO) mice to explore this phenomenon. To generate *Fkbp5*^fl/fl^ mice, CRISPR/Cas9-mediated gene editing was used to insert two loxP sequences into exon 3 of the *Fkbp5* gene (Fig. 2A). These *Fkbp5*^fl/fl^ mice were crossed with *Slc1a3*-Cre^ERT^ transgenic mice, resulting in the generation of *Slc1a3*-Cre^ERT^;*Fkbp5*^fl/fl^ mice. No significant differences in general appearance were observed between the *Fkbp5*^fl/fl^ and *Slc1a3*-Cre^ERT^;*Fkbp5*^fl/fl^ groups (Fig. 2B). Genotyping confirmed the presence of the correct genotype; the primers were designed to detect the *Slc1a3*-Cre^ERT^ and *Fkbp5*^fl/fl^ alleles in the DNA of these mice. Tamoxifen (TAM) injections (100 mg/kg) were used to induce a*Fkbp5*-cKO mice (Fig. 2C). Body weight measurements taken before and after TAM induction revealed no differences between *Fkbp5*^fl/fl^ and a*Fkbp5*-cKO mice (Fig. 2D, E). To validate the efficiency of astrocyte-specific knockout, we used an anti-ACSA-2 antibody and magnetic beads to separate ACSA-2-positive astrocytes from ACSA-2-negative cells in the mouse brain. Western blot analysis of FKBP51 expression in sorted ACSA-2-positive astrocytes revealed an average 42% decrease in FKBP51 levels in a*Fkbp5*-cKO mice compared with *Fkbp5*^fl/fl^ mice. GLAST expression, which was used as a control for successful astrocyte sorting, was not significantly different between the two groups (Fig. 2F, G). We then assessed locomotor activity and anxiety-like behavior using the open field test (OFT) and *Fkbp5*^fl/fl^ and a*Fkbp5*-cKO mice and compared the results with those for WT and *Fkbp5*-KO mice. There were no significant differences in locomotor activity or anxiety levels between the *Fkbp5*^fl/fl^ and a*Fkbp5*-cKO mice or between the WT and *Fkbp5*-KO mice (Fig. 2H, I).

**Fig. 2.**
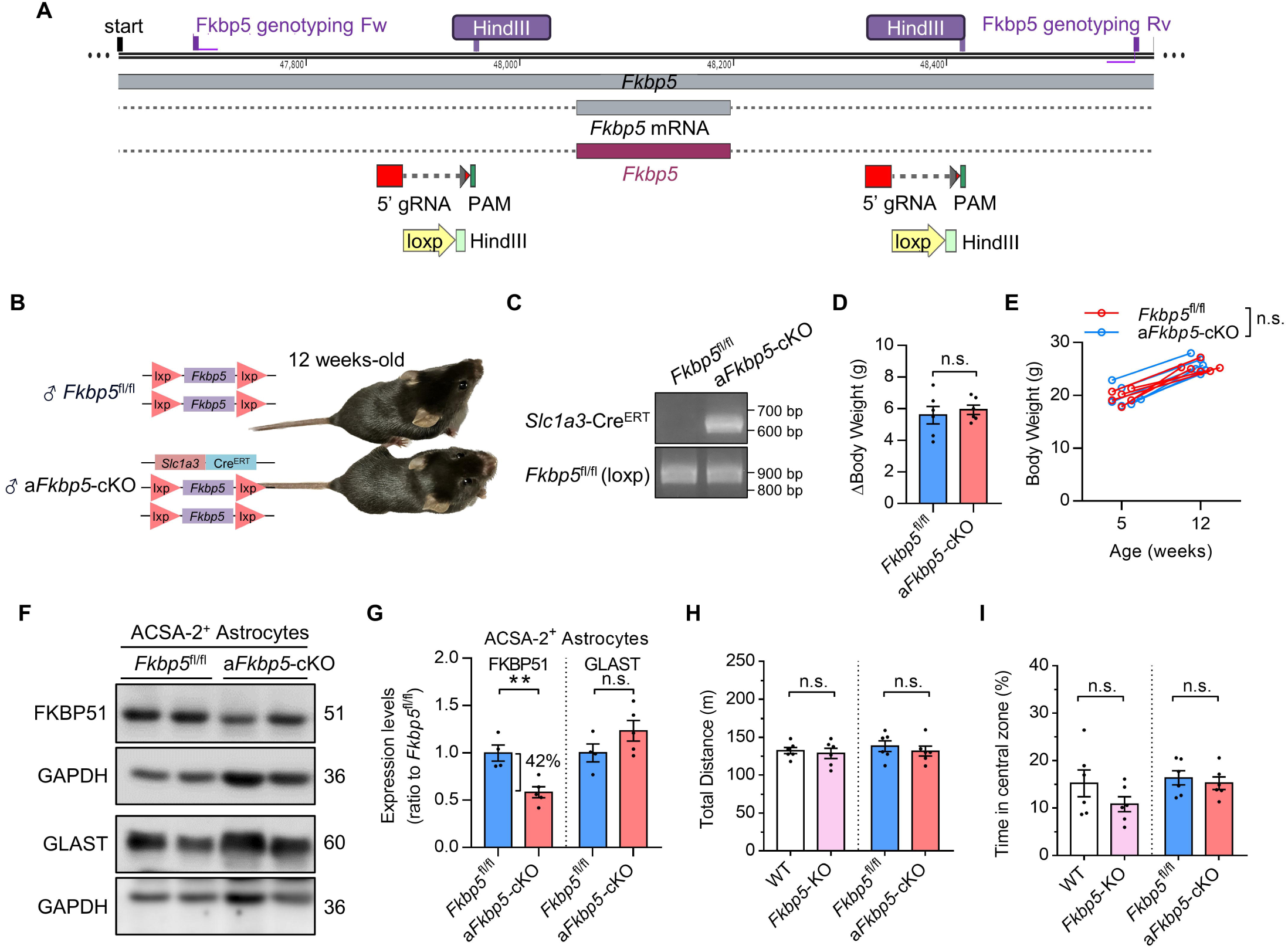
Establishment and characterization of inducible astrocyte-specific *Fkbp5* conditional KO mice. (**A**) CRISPR strategy used to generate the *Fkbp5*^fl/fl^ allele. (**B**) Genotype and appearance of 12-week-old *Fkbp5*^fl/fl^ or astrocytic *Fkbp5* conditional KO (a*Fkbp5*-cKO, *Slc1a3*-Cre^ERT^;*Fkbp5*^fl/fl^) mice. (**C**) Genotyping results confirming *Slc1a3*-Cre^ERT^ and *Fkbp5*^fl/fl^ (loxP) alleles. (**D, E**) Body weight was measured before and 7 weeks after Tamoxifen induction (100 mg/kg/day, 5 days; *n* = 6 per group). (**F, G**) Western blot analysis of the expression of FKBP51 and GLAST in sorted ACSA-2-positived astrocytes from the brains of *Fkbp5*^fl/fl^ or a*Fkbp5*-cKO mice 7 weeks after induction with Tamoxifen. GLAST expression was evaluated to confirm the astrocytic fraction obtained through sorting. *Fkbp5*^fl/fl^ *n* = 4, a*Fkbp5*-cKO *n* = 5. (**H, I**) Open field test used to assess locomotor activity (H, total distance traveled) and anxiety levels (I, time spent in the central zone; lower percentage indicates higher anxiety) in WT and *Fkbp5*-KO mice and in *Fkbp5*^fl/fl^ and a*Fkbp5*-cKO mice (*n* = 6 per group). Data are expressed as the mean ± SEM. In D, G, H, and i, \**P* < 0.05 and \*\**P* < 0.01 for comparisons between groups using an unpaired Student’s *t* test. In E, no significant difference was observed between the two groups using two-way ANOVA followed by Tukey’s post hoc test.

### 3.3. Astrocytic Fkbp5 deletion attenuates KA-induced seizures, cognitive impairment, and hippocampal GLT-1 loss

We investigated the role of FKBP51 in astrocyte-mediated excitotoxicity using a systemic KA-induced seizure model. For a more cost-effective way to examine TAM-induced aFkbp5-cKO mice, we switched to intraperitoneal KA (KA-ip) administration because this modelling approach not only reliably induces seizures and astrogliosis but also has a higher survival rate than does KA-icv by day 7 according to our previous study (Kuo et al., 2019), providing an optimal timeline for assessing FKBP51 function in KA-induced seizures and astrogliosis (Fig. 3A). After KA-ip, a*Fkbp5*-cKO mice exhibited greater resistance to KA-induced seizures than did *Fkbp5*^fl/fl^ control mice (Fig. 3B). Additionally, in the novel object recognition (NOR) test (Kuo et al., 2019), a*Fkbp5*-cKO mice did not display cognitive impairment, in contrast to *Fkbp5*^fl/fl^ control mice (Fig. 3C, D). These results strongly indicate a critical role for astrocytic FKBP51 in KA-induced seizures and cognitive impairment. Next, we investigated astrogliosis in the hippocampus using Western blot analysis. Compared with *Fkbp5*^fl/fl^ control mice*, akbp5*-cKO mice presented significantly lower GFAP levels (Fig. 3E, F). Consistent with these results, a*Fkbp5*-cKO mice presented attenuated GFAP upregulation, specifically in the hippocampal CA3 region (Fig. 3G, H). Hippocampal glutamate transporter-1 (GLT-1), which is mainly expressed on the perisynaptic astrocytic processes and sensitive to excitotoxic injury, particularly in the CA3 region^29^, did not differ in total GLT-1 levels among the groups (Fig. 3E, F). Immunostaining revealed the loss of GLT-1 in the CA3 regions in *Fkbp5*^fl/fl^ control mice but not in a*Fkbp5*-cKO mice, especially in the stratum pyramidale (SP), stratum lucidum (SL), and stratum radiatum (SR) regions. (Fig. 3I, J). Together, these findings suggest that astrocytic *Fkbp5* deletion attenuates KA-induced seizures, cognitive impairment, and hippocampal astrogliosis, while preserving GLT-1 expression in vulnerable CA3 subregions. These results highlight astrocytic FKBP51 as a critical mediator of excitotoxic brain injury.

**Fig. 3.**
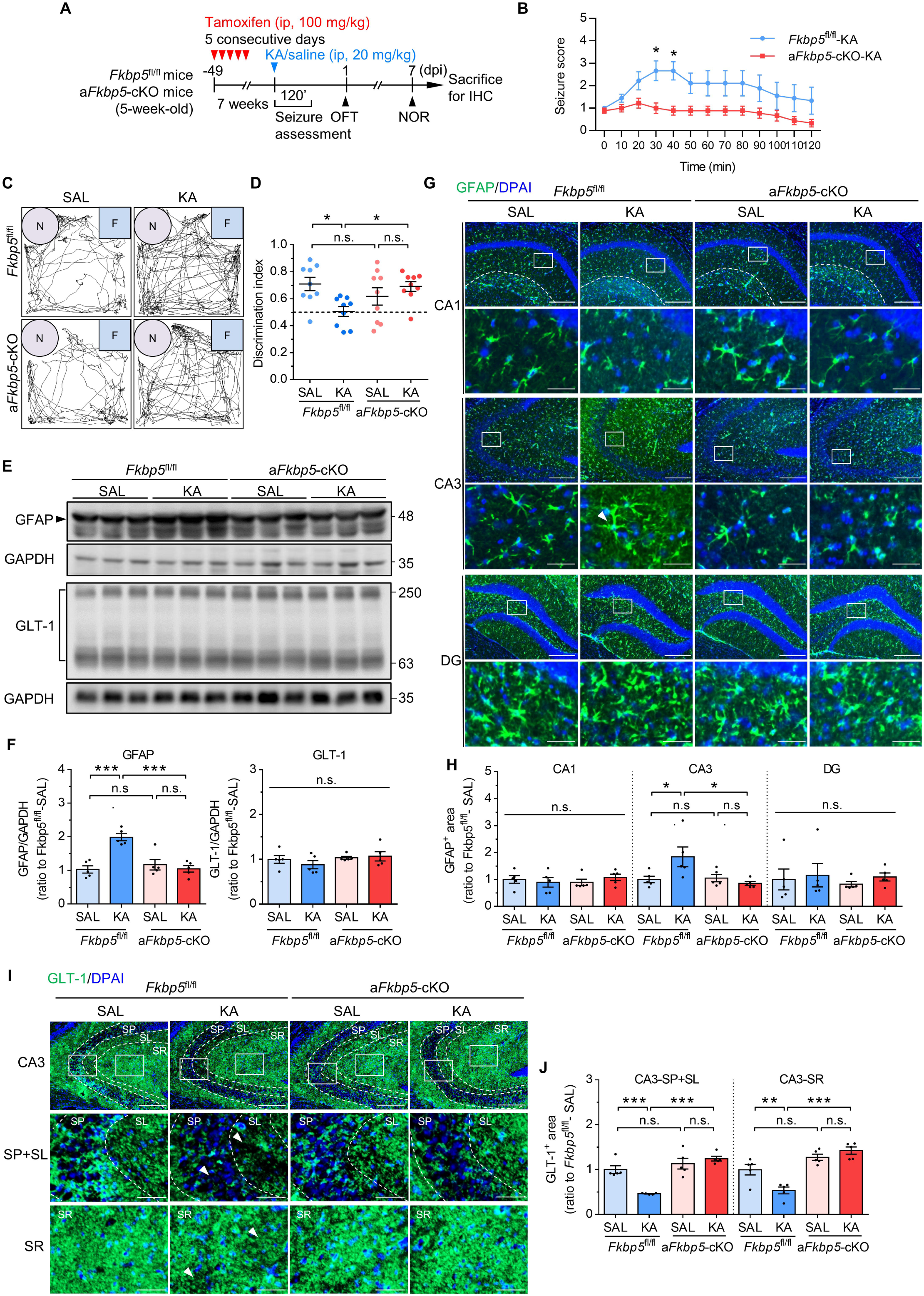
Effects of the conditional KO of astrocytic *Fkbp5* on KA-induced seizures, cognitive impairment, and astrogliosis. (**A**) Scheme of the experimental design. Twelve-week-old *Fkbp5*^fl/fl^ or a*Fkbp5*-cKO mice were subjected to behavioral tests and tissue collection on the 7th day after a single intraperitoneal (ip) injection of saline or KA (3 mg/kg). (**B**) Seizure scores were tracked for 120 minutes after KA injection (*n* = 9 mice per group). \**p* < 0.05 for the comparison between two groups in a two-way repeated measures ANOVA followed by Bonferroni’s post hoc test. (**C**) Representative traces from the novel object recognition (NOR) test for each experimental condition. (**D**) Discrimination index in the NOR test (*n* = 9 mice per group). (**E, F**) Western blot analysis of GFAP and GLT-1 expression in the hippocampi of *Fkbp5*^fl/fl^ and a*Fkbp5*-cKO mice 7 days after the ip injection of saline or KA. Protein levels were normalized to GAPDH (*n* = 5 per group). (**G**) Representative immunofluorescence images of GFAP (green) counterstained with DAPI (blue) in the hippocampal CA1 (upper panel), CA3 (middle panel), and DG (lower panel) regions (scale bar: 200 μm), with a zoomed-in view of the indicated region (scale bar: 50 μm). Arrowhead: hypertrophic astrocyte. (**H**) GFAP area was quantified as the percentage of the immunoreactive area in each hippocampal region of interest and then normalized to that in the *Fkbp5*^fl/fl^ saline treatment group (*n* = 6 per group). (**I**) Representative immunofluorescence images of GLT-1 (green) counterstained with DAPI (blue) in hippocampal CA3 subregions (scale bar: 200 μm), with a zoomed-in view of the stratum pyramidale (SP), stratum lucidum (SL), and stratum radiatum (SR) layers (scale bar: 50 μm). Arrowhead: decreased GLT-1 area. (**J**) The GLT-1 area was quantified as the percentage of the immunoreactive area in each hippocampal region of interest in the SP+SL and SR layers and then normalized to that in the *Fkbp5*^fl/fl^ saline group (*n* = 6 per group). The data are expressed as the mean ± SEM. \**P* < 0.05, \*\**P* < 0.01, and \*\*\**P* < 0.001 for comparisons between two groups using two-way ANOVA followed by Tukey’s post hoc test.

### 3.4. Fkbp5 deletion alleviates excitotoxin-induced neurotoxicity and astrogliosis via NF-***κ***B signaling in primary glia–neuron mixed cultures

Given the in vivo findings, we subsequently explored the mechanisms of FKBP51-dependent astrocyte reactivation using an in vitro glia–neuron (GN) mixed culture system. Immunocytochemistry revealed that excitotoxin NMDA treatment damaged microtubule-associated protein-2 (MAP2)-positive neurons and increased the number of GFAP-positive astrocytes in WT GN cultures (Fig. 4A, B, C). In contrast, in *Fkbp5*-KO GN cultures, such NMDA-induced neurotoxicity and astrogliosis were mitigated (Fig. 4A, B, C), indicating that FKBP51 is pivotal in mediating excitotoxin-induced neurotoxicity and astrogliosis. Next, we investigated which FKBP51-mediated signaling pathways were involved in GN mixed cultures, including the NF-κB (pleiotropic effects in neurons and glial cells, such as cell survival and immune responses) (Dresselhaus and Meffert, 2019) and PH domain leucine-rich repeat-containing protein phosphatase 1 (PHLPP1)-AKT signaling pathways (survival effects against NMDA neurotoxicity) (Wang et al., 2013). Western blot analysis revealed that NMDA treatment did not alter FKBP51 expression (Fig. 4D, E), a finding that aligns with the in vivo hippocampal results (Fig. 1G, K). NMDA treatment led to a reduction in PHLPP1 expression and increased the phosphorylation of p65 and AKT (Fig. 4D, E, F, G, H). Notably, the level of NMDA-induced p65 phosphorylation, a key indicator of NF-κB activation, was significantly decreased in *Fkbp5*-KO GN cultures (Fig. 4D, F), suggesting impaired NF-κB signaling in the absence of FKBP51. PHLPP1 depletion was also detected in *Fkbp5*-KO GN cultures; however, whereas NMDA-induced AKT phosphorylation were not difference between WT and *Fkbp5*-KO GN cultures (Fig. 4D, G, H). These results suggest that *Fkbp5* deletion alleviates NMDA-induced neurotoxicity and astrogliosis, possibly by suppressing NF-κB signaling rather than the PHLPP1-AKT pathway.

**Fig. 4.**
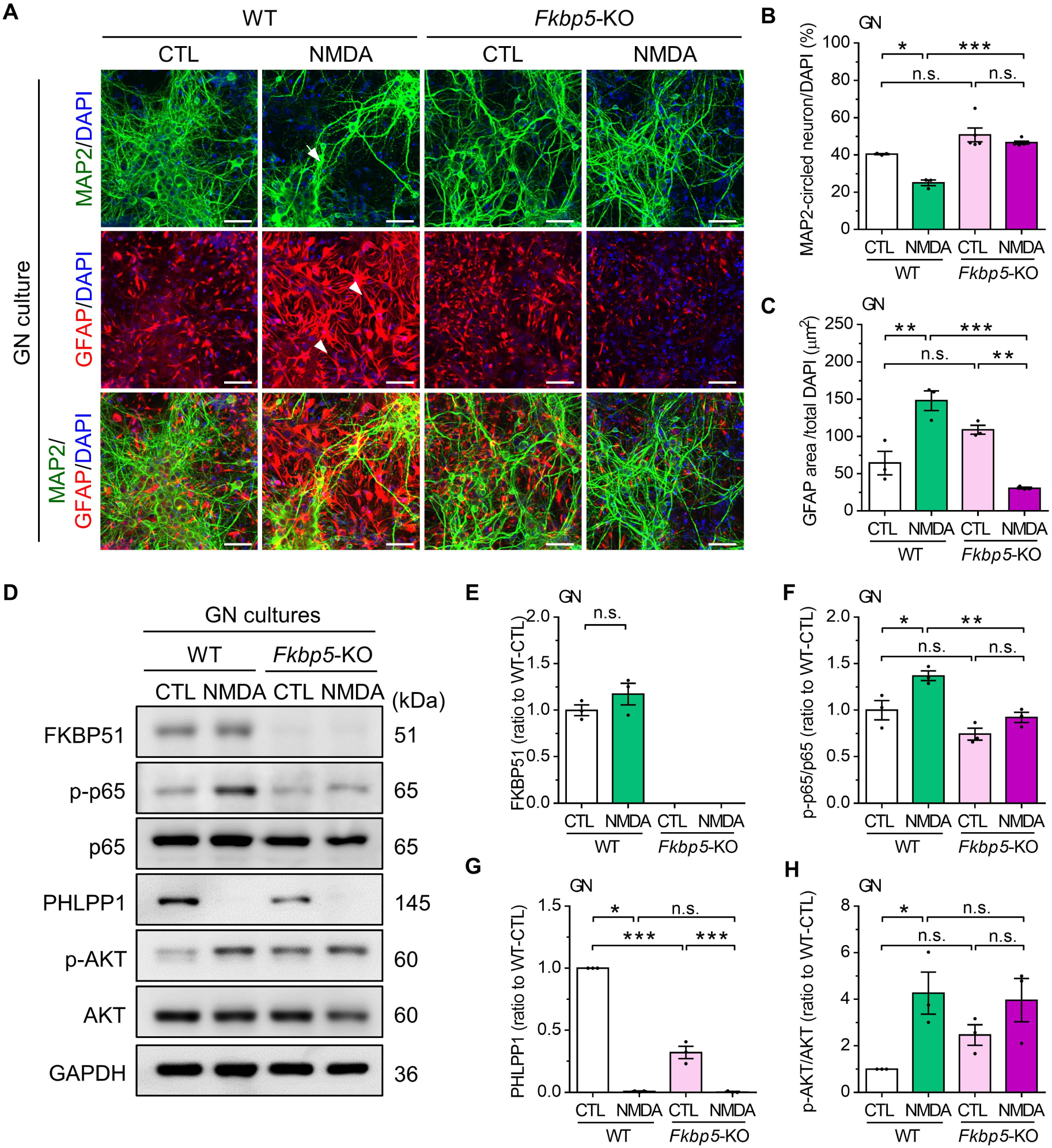
*Fkbp5* deletion reverses NMDA-induced excitotoxicity and the NF-κB signaling pathway in mixed glia–neuron cultures. Primary WT and *Fkbp5*-KO mixed glia**–**neuron (GN) cultures were treated with NMDA (5 μM) for 24 hours. (**A**) Representative immunofluorescence images of MAP2 (green) and GFAP (red) counterstained with DAPI (blue). Scale bar: 50 μm. Arrow: MAP2-circled neuronal loss. Arrowhead: increased GFPA expression and astrocytic hypertrophy. (**B, C**) Quantification of MAP2-circled neurons per DAPI (WT *n* = 3, *Fkbp5*-KO *n* = 5) and GFAP area per DAPI (*n* = 3 per group). (**D**) Western blot analysis of FKBP51, phosphorylated p65 (p-p65), p65, PHLPP1, phosphorylated AKT (p-AKT), and AKT. (**E**–**H**) Quantification of FKBP51 (E), p-p65/p65 (F), PHLPP1 (G), and pAKT/AKT (H) (*n* = 3 per group). Data are expressed as the mean ± SEM. \**P* < 0.05, \*\**P* < 0.01, and \*\*\**P* < 0.001 for the comparison between two groups by two-way ANOVA followed by Tukey’s post hoc test.

### 3.5. Disruption of the FKBP51–IKK***α*** interaction by Fkbp5-3AR mutants reduces lipopolysaccharide (LPS)-induced NF-***κ***B p65 phosphorylation in astrocytes

On the basis of the observation of FKBP51-dependent NF-κB signaling in excitotoxicity and astrogliosis, we next employed computer modeling to explore the intricate interaction between FKBP51 and NF-κB in primary astrocyte cultures. Docking with each energy parameter set resulted in ten modeling candidates defined by the centers of highly populated clusters of low-energy docked structures. Among them, the FKBP51/IKKα model suggested that FKBP51 and IKKα formed a long groove that might be suitable for NF-κB/IκB binding, thereby facilitating the kinase activity of IKKα. Given that FKBP51 is known to promote IκB degradation by binding to IKKα (Tong and Jiang, 2015), we designed a Myc-tagged *Fkbp5* quadruple mutant (Myc-*Fkbp5*-3AR: E258A, E301A, Y302A, and G345R) to interfere with FKBP51 and IKKα interactions to investigate the effects of FKBP51 on NF**-**κB activation and the inflammatory response. Primary mouse astrocytes were transfected with either a Myc vector, Myc-tagged wild-type *Fkbp5* (Myc-*Fkbp5*-WT), or the Myc-*Fkbp5*-3AR mutant, resulting in the robust overexpression of *Fkbp5* mRNA in astrocytes (Fig. 5A). To simulate neuroinflammation and NF-κB signaling in astrocytes, the cells were treated with the TLR4 agonist LPS, a well-established activator of NF-κB in astrocytes, to assess the effects of the *Fkbp5* mutant on NF-κB-driven inflammatory responses. Western blot analysis confirmed similar levels of Myc-tagged *Fkbp5* between astrocytes transfected with Myc-*Fkbp5*-WT and those transfected with Myc-*Fkbp5*-3AR (Fig. 5B, C). No changes in FKBP51 levels were observed after LPS treatment (Fig. 5B, D). Importantly, LPS treatment significantly increased p65 phosphorylation in both the Myc-vector and Myc-*Fkbp5*-WT groups. In contrast, compared with Myc-*Fkbp5*-WT, the Myc-*Fkbp5*-3AR mutant markedly decreased LPS-induced p65 phosphorylation (Fig. 5B, E), suggesting that the Myc-*Fkbp5*-3AR mutant effectively suppressed NF-κB activation. FKBP51 is known to regulate the expression of NF-κB-driven inflammatory genes, including *Tnf-*α and *Il-1*β (Gan et al., 2024; Kästle et al., 2018). Reverse transcription–quantitative PCR (RTLqPCR) analysis revealed a significant increase in *Tnf-*α and *Il-1*β mRNA expression after LPS treatment. Notably, the Myc-*Fkbp5*-3AR mutant significantly reduced LPS-induced *Tnf-*α gene expression but did not affect *Il-1*β expression in astrocytes (Fig. 5F, G). Although *Fkbp5* is also known to regulate the AKT pathway by interacting with PHLPP1 (Tong and Jiang, 2015), we found that AKT phosphorylation was unaffected by the Myc-*Fkbp5*-3AR mutation. Our results indicate that this mutant specifically targets FKBP51-NF-κB signaling and inhibits p65 phosphorylation without affecting AKT (Fig. 5H, I) in inflammatory astrogliosis.

**Fig. 5.**
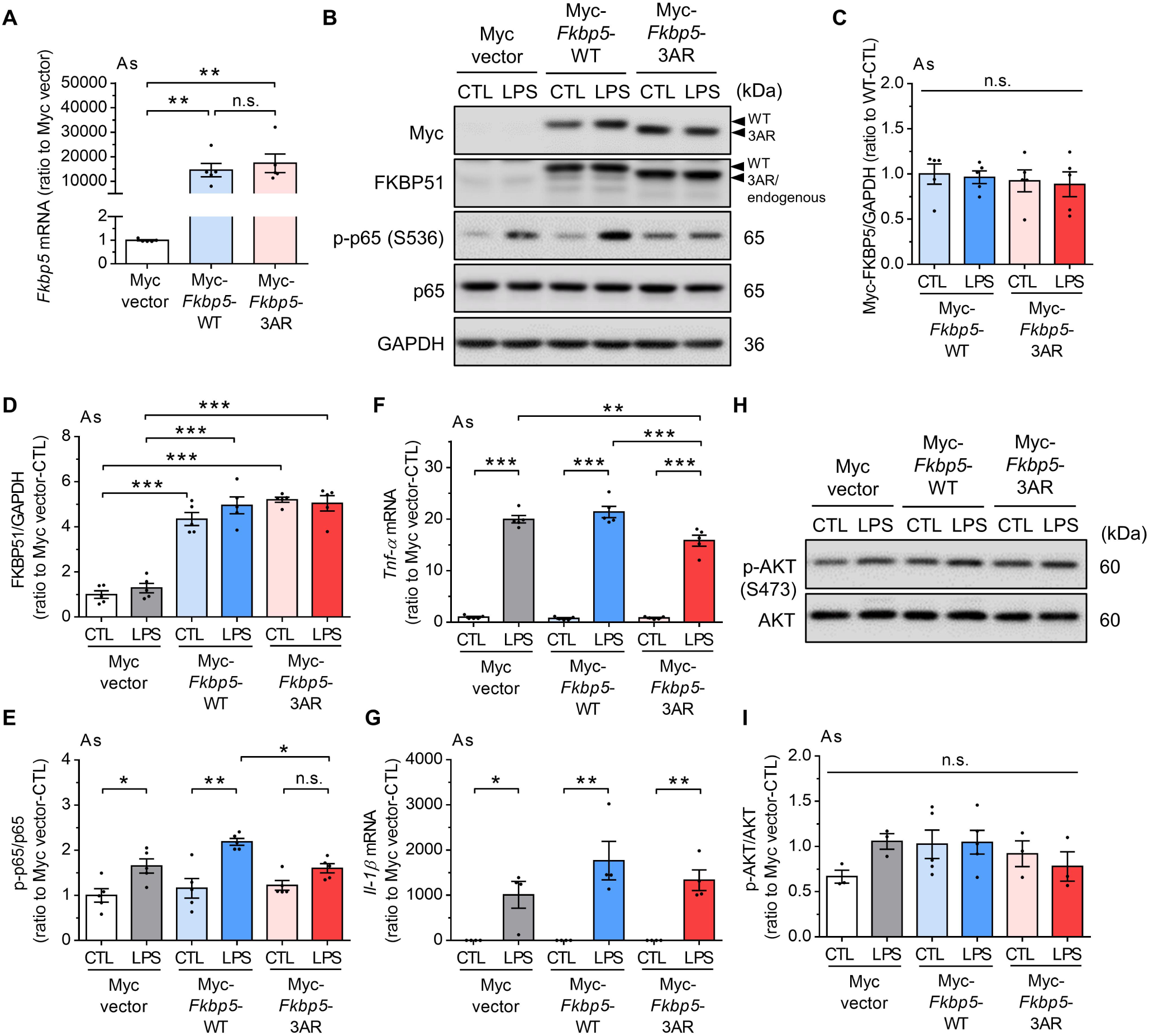
The *Fkbp5*-3AR mutant specifically decreases LPS-induced NF-κB activation in cultured astrocytes. (**A**L**E**) Primary mouse astrocytes were transfected with Myc-vector, Myc-*Fkbp5*-WT or Myc-*Fkbp5*-3AR for 24 hours and treated with 1 μg/ml LPS for 1 hour, followed by Western blot analysis of Myc, FKBP51, phospho-p65 (p-p65), and p65. The phosphorylated signals were quantified as the ratio of phosphorylated protein to total protein levels and normalized to the level of GAPDH (*n* = 5 per group). (**F, G**) RTLqPCR analysis of *Tnf-*α and *Il-1*β mRNA (*n* = 5 per group). The housekeeping gene 18S was used as the internal control. (**H**) Western blot analysis of phospho-AKT (p-AKT) and AKT. (**I**) The phosphorylated signals were quantified as the ratio of phosphorylated protein to total protein levels and normalized to the level of GAPDH (*n* = 5 per group). The data are expressed as the mean ± SEM. In **I**, \**P* < 0.05 and \*\**P* < 0.01 compared between the two indicated groups by one-way ANOVA with Tukey’s post hoc test. In **C**L**I**, \**P* < 0.05, \*\**P* < 0.01, and \*\*\**P* < 0.001 compared between the two indicated groups by two-way ANOVA with Tukey’s post hoc test.

### 3.6. Hippocampal transcriptomic profiles after LPS treatment revealed the enrichment of NF-***κ***B signaling, which was counteracted by Fkbp5 deletion

To comprehensively examine in vivo gene expression changes associated with neuroinflammation and FKBP51 deletion, we conducted bulk RNA sequencing using hippocampal tissue from WT and *Fkbp5*-KO mice treated with either saline (SAL) or LPS for 7 days. Differential gene expression analysis revealed 195 upregulated and 226 downregulated genes between the *Fkbp5*-KO-LPS and WT-LPS groups and 248 upregulated and 108 downregulated genes between the WT-LPS and WT-SAL groups (Fig. 6A). We conducted gene set enrichment analysis (GSEA) using the Hallmark gene sets, and the results revealed a high normalized enrichment score (NES) for inflammatory-related pathways, including interferon responses, for the WT-LPS vs. WT-SAL comparison (Fig. 6B). In contrast, for the *Fkbp5*-KO-LPS vs. WT-LPS comparison, the NES was low for the interferon alpha response but high for glycolysis, a metabolic process linked to astrocyte function in the brain (Fig. 6C). To investigate the biological processes involved, we performed GSEA using the Gene Ontology (GO) biological process database. The results highlighted enrichment in glutamate-related signaling, glial cell-associated processes, inflammatory responses, and NF-κB signaling. Notably, most of these pathways were activated in the WT-LPS vs. WT-SAL comparison but counteracted in the *Fkbp5*-KO-LPS vs. WT-LPS comparison (Fig. 6D). The individual enrichment plots further revealed that the “inflammatory response” was strongly activated in the WT-LPS vs. WT-SAL comparison (NES: 1.96; *p* < 0.001) but significantly suppressed in the *Fkbp5*-KO-LPS vs. WT-LPS comparison (NES: -1.28; *p* < 0.05) (Fig. 6E). Similarly, “canonical NF-κB signal transduction” was activated in the WT-LPS vs. WT-SAL comparison (NES: 1.66; *p* < 0.001) but suppressed in the *Fkbp5*-KO-LPS vs. WT-LPS comparison (NES: -1.35; *p* < 0.05) (Fig. 6E), indicating that FKBP51 mediates NF-κB signaling and is involved in inflammatory processes. Ingenuity pathway analysis (IPA) of DEGs between the *Fkbp5*-KO-LPS and WT-LPS groups revealed several affected upstream regulators, including NFKB1, SOX2, ETV1, TCF7L2, and NFE2L2. Notably, NFKB1 was significantly inhibited, further supporting the suppression of NF-κB signaling in *Fkbp5*-KO mice *Fkbp5* (Fig. 6F). Collectively, these results underscore the pivotal role of FKBP51 in driving NF-κB-mediated neuroinflammation.

**Fig. 6.**
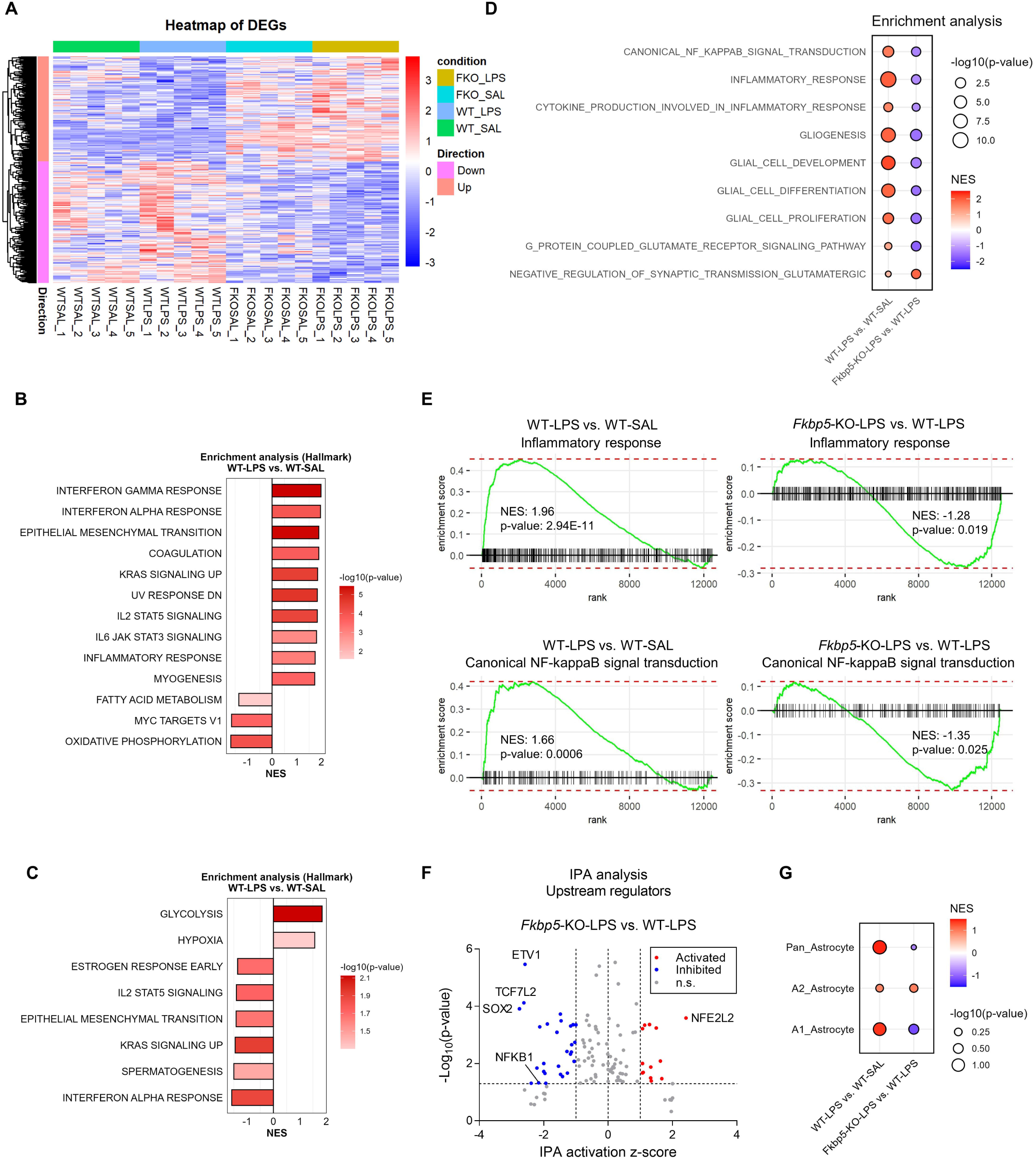
Transcriptional profile analysis of the hippocampus of *Fkbp5*-KO mice in an inflammatory state. (**A**) Bulk RNA sequencing was performed using hippocampi from WT and *Fkbp5*-KO mice 7 days after LPS injection (3 mg/kg; *n* = 5 per group). Differentially expressed genes (DEGs) were identified by comparing the *Fkbp5*-KO-LPS and WT-LPS groups (195 upregulated and 226 downregulated) and WT-LPS and WT-SAL groups (248 upregulated and 108 downregulated). The direction of gene expression change was defined based on the *Fkbp5*-KO-LPS vs. WT-LPS comparison. (**B, C**) Gene set enrichment analysis (GSEA) using the Hallmark gene sets for the WT-LPS and WT-SAL groups and the *Fkbp5*-KO-LPS v and WT-LPS groups. The color scale represents the -log10(*P* value). (**D**) GSEA using the Gene Ontology Biological Processes database revealed significant enrichment in glutamate signaling, glial-related processes, and inflammatory responses, including NF-κB signaling. These processes were inversely regulated in the two comparisons: activated in the WT-LPS vs. WT-SAL comparison and suppressed in the *Fkbp5*-KO-LPS vs. WT-LPS comparison. The color scale represents the normalized enrichment score (NES), and the dot size reflects -log10(*P* value). (**E**) Enrichment plots revealed that inflammatory responses (upper panel) and canonical NF-κB signal transduction (lower panel) were activated in the WT-LPS vs. WT-CTL comparison but inhibited in the *Fkbp5*-KO-LPS vs. WT-LPS comparison. (**F**) Volcano plot displaying the IPA activation Z score on the x-axis and - log10(*P* value) on the y-axis. The dashed lines on the x-axis represent Z scores of 1 and -1, whereas the dashed line on the y-axis indicates a *P* value threshold of 0.05. The blue dots represent inhibited regulators, and the red dots indicate activated regulators. IPA revealed key upstream regulators, highlighting significant NF-κB inhibition in *Fkbp5*-KO-LPS mice compared with WT-LPS mice. (**G**) GSEA of astrocyte subtype gene signatures (panreactive, A1, and A2) revealed trends toward increased panreactive and A1 markers in the WT-LPS vs. WT-SAL comparison and a reduction in A1 markers in the *Fkbp5*-KO-LPS vs. WT-LPS comparison.

Given that FKBP5 has been suggested as an A1-type proinflammatory astrocyte marker, we evaluated astrocyte subtype signatures using GSEA. Although no significant enrichment was observed in the panreactive, A1, or A2 astrocyte signatures, there was a trend toward increased NESs for panreactive and A1 astrocyte markers in the WT-LPS vs. WT-SAL comparison and a mild reduction in A1 markers in the *Fkbp5*-KO-LPS vs. WT-LPS comparison (Fig. 6G). These results support the role of FKBP51 in driving NF-κB-mediated neuroinflammation, glial function, and glutamatergic signaling.

## 4. Discussion

The role of the FKBP51 cochaperone in regulating stress responses through the modulation of GR sensitivity in the HPA axis is well established, linking it to stress-related psychiatric disorders (Binder, 2009; Hartmann et al., 2015; Zannas et al., 2016). In addition to interacting with GR, FKBP51 engages with other signaling pathways, notably IKKα/NF-κB and AKT/PHLPP1, in a range of cellular functions (Romano et al., 2011; Tufano et al., 2023). Recent studies have suggested that FKBP51 is also involved in metabolic functions, aging, neuropathic pain, and even acute ischemic stroke (Balsevich et al., 2017; Hartmann et al., 2012; Maiarù et al., 2016; Yu et al., 2020). In this study, we provide the first evidence that, in excitotoxin KA-induced seizure models, FKBP51 plays a pivotal role in activating astrogliosis via NF-κB signaling, thus providing a therapeutic target against excitotoxic brain injury. Our results revealed that compared with WT control mice, complete *Fkbp5*-KO mice presented fewer KA-induced seizures and less neuronal loss and astrogliosis. Importantly, in established astrocyte-specific conditional a*Fkbp5*-cKO mice, we observed a similar reduction in KA-induced seizures and astrogliosis, along with improvements in novel object recognition ability. The preservation of glutamate transporter-1 (GLT-1) levels only in the hippocampal CA3 region of a*Fkbp5*-cKO mice underscores that astrocyte-specific FKBP51 ablation maintains glutamate homeostasis against excitotoxicity. Furthermore, in vitro glia–neuron mixed cultures from *Fkbp5*-KO mice ameliorated NMDA-induced neurotoxicity and astrogliosis, accompanied by a decrease in the phosphorylation of the NF-κB p65 subunit. Our manufactured *Fkbp5*-3AR mutant successfully interrupted the FKBP51-NF-κB interaction in primary cultures, a finding that supports the targeting of astrocytic FKBP51 function in NF-κB signaling. These findings suggest that FKBP51 is a critical regulator of proinflammatory astrogliosis and excitotoxicity mediated through NF-κB signaling. The reduced NF-κB activation in response to FKBP51 ablation highlights the importance of this pathway in modulating neuroinflammation during excitotoxic stress. Collectively, these results provide compelling evidence that FKBP51 influences astrocyte activation via NF-κB regulation, modulating GLT-1 expression in excitotoxic brain injury, and provide insights into the role of FKBP51 in astrocyte-mediated neuroprotection and excitotoxicity.

FKBP51 has been implicated in regulating synaptic plasticity and neurotransmission, for example, long-term depression (LTD) and long-term potentiation (LTP) (Blair et al., 2019; Qiu et al., 2019). Our previous study, in line with those findings, revealed that *Fkbp5*-KO mice subjected to the potent immune activator LPS presented enhanced anxiety-like behavior, a reduced immune response and microglial activation in the hippocampus. In contrast to the increase in glutamic acid decarboxylase 65 (GAD65) and GABA synthesis in LPS-treated WT mice, GAD65 was not increased in *Fkbp5*-KO mice, suggesting that FKBP5 potentially influences inhibitory neurotransmission under inflammatory conditions (Gan et al., 2022). Previous research on FKBP5 mouse models, as reviewed by Gebru *et al*. (Gebru et al., 2024), has focused primarily on its role in stress regulation and psychiatric disorders. While much of the literature has explored the involvement of FKBP51 in these contexts, our study extends these findings by demonstrating its functional relevance in astrocyte-mediated excitotoxicity and seizure pathophysiology. To bridge this gap, we generated the first astrocyte-specific conditional *Fkbp5* knockout mouse. Notably, we found that FKBP51 ablation in astrocytes led to a decrease in GFAP expression, accompanied by an increase in GLT-1 in the hippocampal CA3 subregion, which is particularly vulnerable to KA-induced damage. The increased expression of GLT-1, a key glutamate transporter, points to an enhanced ability of astrocytes to regulate glutamate clearance, potentially mitigating the excitotoxic effects caused by excessive glutamate, a hallmark of both acute and chronic neurodegenerative conditions (Fontana, 2015; Rothstein et al., 1996). These findings broaden our understanding of the involvement of FKBP51 in neural plasticity and highlight its potential role in mitigating excitotoxic brain injuries.

The interaction of FKBP51 with IKKα/NF-κB signaling is central to the regulation of inflammation and immune responses (Chen and Greene, 2004; Guo et al., 2024; Liu et al., 2017b). Our results suggest the novel concept of FKBP51-mediated NF-κB signaling in astrogliosis in the excitotoxic state in the brain. Recent research has revealed that a proinflammatory astrocyte subpopulation induced by activated microglia contributes to neurotoxicity and neurodegeneration (Clarke et al., 2018; Liddelow et al., 2017). While these studies did not directly implicate FKBP51 as a proinflammatory astrocytic marker, our study extends the understanding of the role of FKBP51 in astrocyte biology. We provide mechanistic insights into how FKBP51 ablation in both neurons and astrocytes enhances resilience to excitotoxic insults, suggesting that FKBP51 is involved in proinflammatory glial responses under pathological conditions. In response to FKBP51 ablation, astrocytes may adopt a more protective phenotype, promoting neuronal survival and synaptic stability under excitotoxic stress. Further research is needed to elucidate the broader effects of FKBP51 modulation in astrocytes, the results of which could provide more insights into how astrocytes respond to excitotoxic injury and other neurological disorders.

For the treatment of epilepsy, most antiseizure medications primarily target ion channels, such as sodium or calcium channels, often leading to side effects such as dizziness, cognitive impairment, and motor incoordination(Asadi-Pooya et al., 2023). These adverse effects arise from the broad influence of ion channel blockade on normal neuronal activity. Moreover, a substantial proportion of epilepsy patients, particularly those with drug-resistant epilepsy, do not respond to these therapies because of complex mechanisms involving changes in transporter activity, alterations in the neuronal network, underlying brain diseases, and genetic variations (Lin et al., 2021). These findings underscore the need for innovative therapeutic strategies beyond ion channel modulation. Clinical and experimental evidence has highlighted the central role of neuroinflammation in epileptogenesis and drug resistance, with astrogliosis playing a critical role in this process (Rana and Musto, 2018; Vezzani et al., 2011; Vezzani et al., 2013). Astrocytes perform numerous physiological functions, including synapse formation, ion homeostasis, neurotransmitter regulation, blood brain barrier component, and neuroinflammation. Our findings demonstrate that astrocytic *Fkbp5* deletion is promising for reducing KA-induced seizure activity and NF-κB signaling. Targeting the FKBP51-NF-κB axis represents a promising non-ion channel therapeutic strategy. The results from our docking model and FKBP51-3AR mutant experiments support the concept of targeting astrocytic FKBP51 to regulate NF-κB signaling. This approach may further attenuate seizures as well as neuroinflammation without directly influencing ion channel function, potentially minimizing cognitive and motor impairment.

We did not perform a transcriptomic analysis of astrocyte-specific a*Fkbp5*-cKO mice to examine NF-κB signaling under excitotoxic conditions, which is a limitation of this study. This approach would have provided insights into the role of astrocytic FKBP51 in excitotoxicity. Future studies employing single-cell RNA sequencing or spatial transcriptomics in a*Fkbp5*-cKO mice can dissect the transcriptional changes in astrocyte subpopulations with regional specificity, particularly in the CA3 subregion, where FKBP51 ablation is associated with upregulated GLT-1 expression and neuroprotection. Such findings would help to elucidate the astrocyte-mediated mechanisms of FKBP51 in excitotoxic injury and neuroinflammation, providing a more detailed understanding of the role of FKBP51 at the cellular and circuit levels. Moreover, our *Fkbp5*-3AR mutant was designed to specifically disrupt the interaction of FKBP51 and IKKα, and we demonstrated that it reduced the LPS-induced phosphorylation of p65. However, further studies are needed to confirm whether the *Fkbp5*-3AR mutant selectively affects FKBP51-IKKα binding without altering interactions with other FKBP51-targeted molecules. This specificity is essential for defining the mechanism of FKBP51-mediated NF-κB signaling regulation.

## 5. Conclusions

The current study demonstrated that astrocytic FKBP51 ablation leads to reduced excitotoxicity, seizure activity, and astrogliosis, highlighting the preservation of glutamate homeostasis via astrocytic GLT-1 and the modulation of NF-κB signaling as critical mechanisms through which FKBP51 influences brain functions. Specific interruption of FKBP5-IKKα interaction presents an opportunity to reshape strategies for treating neuroinflammation and excitotoxicity.

## Abbreviations

a*Fkbp5*-cKO: Astrocyte-specific *Fkbp5* conditional knockout
AKT: Protein kinase B
CNS: Central nervous system
FKBP51: FK506-binding protein 51
*Fkbp5*-KO: *Fkbp5* knockout
GLT-1: Glutamate transporter-1
GN: glia-neuron
GR: Glucocorticoid receptor
HPA: Hypothalamic–pituitary–adrenal
HPC: Hippocampus
IKK: IκB kinase
KA: Kainic acid
LPS: Lipopolysaccharide
NF-κB: Nuclear factor kappa-light chain enhancer of activated B cells
NMDA: N-methyl-D-aspartate
OL: Oligodendrocyte
PHLPP1: PH domain leucine-rich repeat-containing protein phosphatase 1
RTLqPCR: Reverse transcription quantitative polymerase chain reaction
SAL: Saline
WT: Wild-type

## Ethics approval and consent to participate

All animal experiments were reviewed and approved by the Institutional Animal Care and Use Committee (IACUC) of National Yang Ming Chiao Tung University (IACUC number: 1090515 and 1100336) and performed in accordance with the Guide for the Care and Use of Laboratory Animals by the National Institute of Health.

## Data availability

The data that support the findings in this study are available from the corresponding author upon reasonable request.

## CRediT authorship contribution statement

**Yu-Ling Gan:** Conceptualization, Data curation, Formal analysis, Investigation, Validation, Visualization, Writing – original draft, Writing – review & editing. **Shang-Hsuan Lin:** Data curation, Investigation, Validation, Visualization, Writing – original draft. **Yu-Ping Kang:** Data curation, Investigation, Validation, Visualization, Writing – original draft. **Jia-Zhen Zhou:** Data curation, Investigation, Validation, Visualization. **Wei-Hsuan Huang:** Data curation, Investigation, Validation, Visualization. **Pin-Hua Sung:** Data curation, Investigation, Validation, Visualization. **Chia-Chi Hung:** Data curation, Validation, Visualization, Writing – review & editing. **Pei-Chien Hsu:** Data curation, Validation, Visualization. **Shu-Yin Chu:** Writing – review & editing. **Feng-Shiun Shie:** Writing – review & editing. **I-Hui Lee:** Funding acquisition, Supervision, Writing – review & editing. **Chung-Jiuan Jeng:** Conceptualization, Funding acquisition, Supervision, Writing – review & editing. **Yi-Hsuan Lee:** Conceptualization, Data curation, Formal analysis, Funding acquisition, Project administration, Supervision, Writing – original draft, Writing – review & editing.

## Acknowledgements

We would like to acknowledge the contribution of Dr. Chia-Cheng Chou, who sadly passed away before this work could be completed. His invaluable insights and dedication to this research will always be remembered. We acknowledge the Transgenic Core Facility at the Institute of Molecular Biology, Academia Sinica, Taiwan, for their support in generating *Fkbp5*^fl/fl^ mice using CRISPR technology. We also thank the Biomedical Resource Core at the First Core Labs, National Taiwan University College of Medicine, for their technical assistance in constructing the mutant.

