## Supplementary for "Astrocytic FKBP5 Regulates Neuroinflammation and Cognitive Outcomes in Excitotoxic Brain Injury"

### Supplementary data

**Supplementary Table S1.** Primer sequences used for genotyping.

| Gene | Primer sequence (5' to 3') | Product length |
|---|---|---|
| *Fkbp5-loxP* | Forward: GCCTTGGTGGAGATAGGGTTTCA | 882 bp |
| | Reverse: GTGGACAGAATTAGGTTTACATCTGCC | (*Fkbp5*$^{fl/fl}$ allele) |
| *Slc1a3-Cre*$^{ERT}$ | Forward: ACAATCTGGCCTGCTACCAAAGC | 600 bp |
| | Reverse: CCAGTGAAACAGCATTGCTGTC | (*Slc1a3*-Cre$^{ERT}$ allele) |

**Supplementary Table S2.** Antibodies used for Western blot and immunofluorescence staining.

| Target | Host | Supplier | Cat. No. | Application |
|---|---|---|---|---|
| ACSA-2 | Mouse | Miltenyi Biotec | 130-117-535 | MICS |
| AKT | Rabbit | Cell Signaling Technology | 4691 | WB |
| p-AKT (S473) | Rabbit | Cell Signaling Technology | 9271 | WB |
| Caspr | Mouse | NeuroMab | 75-001 | IF |
| FKBP51 | Rabbit | Cell Signaling Technology | 12210 | WB |
| GAPDH | Rabbit | GeneTex | GTX100118 | WB |
| GFAP | Rabbit | Dako | Z0334 | WB and IF |
| GFAP | Mouse | Merck Millipore | MAB360 | IF |
| GLAST | Rabbit | Abcam | ab416 | WB and IF |
| GLT-1 | Rabbit | Abcam | ab41621 | WB and IF |
| MAP2 | Mouse | Merck Millipore | MAB3418 | IF |
| MBP | Rat | Bio-Rad | MCA409S | WB and IF |
| Myc (Clone 9E10) | Mouse | N/A | N/A | WB |
| $Na_V1.6$ | Rabbit | Alomone Labs | ASC-009 | IF |
| NeuN | Rabbit | Abcam | ab177487 | IF |
| p65 | Rabbit | Cell Signaling Technology | 8242 | WB |
| p-p65 (S536) | Rabbit | Cell Signaling Technology | 3033 | WB |
| PHLPP1 | Rabbit | Merck Millipore | 07-1341 | WB |
| Rip | Mouse | Merck Millipore | MAB1580 | IF |
| HRP-Secondary | — | Jackson ImmunoResearch | | WB |
| anti-rabbit IgG | | Laboratories | 111-035-003 | |
| anti-mouse IgG | | | 115-035-003 | |
| Alexa Fluor-Secondary | — | Invitrogen | | IF |
| anti-mouse IgG, Alexa 488 | | | A21202 | |
| anti-mouse IgG, Alexa 594 | | | A21203 | |
| anti-rabbit IgG, Alexa 488 | | | A21202 | |
| anti-rabbit IgG, Alexa 594 | | | A21207 | |
| anti-rat IgG, Alexa 488 488 | | | A21208 | |

**Supplementary Table S3.** Primer sequences used for RT-qPCR.

| Gene | Primer sequence (5' to 3') | NCBI accession No. |
|---|---|---|
| *Fkbp5* | Forward: AAAGGACAATGACTACTGATGAGG | NM_010220.4 |
| | Reverse: CTGACAACATCCCTTTGTAGTGGAC | |
| *Tnf-α* | Forward: TAGCCCACGTCGTAGCAAAC | NM_013693.3 |
| | Reverse: GCAGCCTTGTCCCTTGAAGA | |
| *Il-1b* | Forward: TGCCACCTTTTGACAGTGATGAG | NM_008361.4 |
| | Reverse: TGCCTGCCTGAAGCTCTTGT | |
| *Mbp* | Forward: CACGAGAACTACCCATTATGGCTCC | NM_001025251.2 |
| | Reverse: GATGGAGGTGGTGTTCGAGGTGT | |
| *Epo* | Forward: CGTAGCCTCACTTCACTGCTTCGG | NM_007942.2 |
| | Reverse: CTCTCCCGTGTACAGCTTCAG | |
| *18S rRNA* | Forward: GAGGTGAAATTTCTTGGACCGG | NR_003278.3 |
| | Reverse: CGAACCTCCGACTTTCGTTCT | |

**Supplementary Table S4.** Summary of statistical analysis.

| Figure | Measurement | Group Comparison | Type of test | Factor | Statistical value | *P* value |
|---|---|---|---|---|---|---|
| **1B** | Seizure Score | WT-KA vs *Fkbp5*-KO-KA | Two-way repeated measures ANOVA with Bonferroni's *post hoc* test | Genotype | F (1, 16) = 11.72 | 0.0035 |
| | | | | Time | F (9, 144) = 34.17 | < 0.0001 |
| | | | | Genotype × Time | F (9, 144) = 1.085 | 0.3774 |
| **1D** | NeuN+ cells, CA1 | WT vs *Fkbp5*-KO | Two-way ANOVA with Tukey's *post hoc* test | Genotype | F (1, 12) = 7.83 | 0.0161 |
| | | | | Treatment | F (1, 12) = 16.8 | 0.0015 |
| | | | | Genotype × Treatment | F (1, 12) = 12.8 | 0.0038 |
| | NeuN+ cells, CA3 | WT vs *Fkbp5*-KO | Two-way ANOVA with Tukey's *post hoc* test | Genotype | F (1, 12) = 3.85 | 0.0735 |
| | | | | Treatment | F (1, 12) = 20.6 | 0.0007 |
| | | | | Genotype × Treatment | F (1, 12) = 3.04 | 0.1069 |
| | NeuN+ cells, DG | WT vs *Fkbp5*-KO | Two-way ANOVA with Tukey's *post hoc* test | Genotype | F (1, 12) = 3.06 | 0.0813 |
| | | | | Treatment | F (1, 12) = 3.32 | 0.0935 |
| | | | | Genotype × Treatment | F (1, 12) = 3.62 | 0.1057 |
| **1F** | GFAP area, CA1 | WT vs *Fkbp5*-KO | Two-way ANOVA with Tukey's *post hoc* test | Genotype | F (1, 8) = 8.79 | 0.0180 |
| | | | | Treatment | F (1, 8) = 11.3 | 0.0098 |
| | | | | Genotype × Treatment | F (1, 8) = 9.87 | 0.0138 |
| | GFAP area, CA3 | WT vs *Fkbp5*-KO | Two-way ANOVA with Tukey's *post hoc* test | Genotype | F (1, 8) = 7.29 | 0.0271 |
| | | | | Treatment | F (1, 8) = 7.86 | 0.0231 |
| | | | | Genotype × Treatment | F (1, 8) = 7.24 | 0.0274 |
| | GFAP area, DG | WT vs *Fkbp5*-KO | Two-way ANOVA with Tukey's *post hoc* test | Genotype | F (1, 8) = 2.80 | 0.1330 |
| | | | | Treatment | F (1, 8) = 2.94 | 0.1248 |
| | | | | Genotype × Treatment | F (1, 8) = 2.61 | 0.1447 |
| **1H** | CC-MBP | WT vs *Fkbp5*-KO | Two-way ANOVA with Tukey's *post hoc* test | Genotype | F (1, 20) = 15.9 | 0.0007 |
| | | | | Treatment | F (1, 20) = 0.0315 | 0.8608 |
| | | | | Genotype × Treatment | F (1, 20) = 21.0 | 0.0002 |
| **1I** | HPC-MBP | WT vs *Fkbp5*-KO | Two-way ANOVA with Tukey's *post hoc* test | Genotype | F (1, 20) = 1.17 | 0.2927 |
| | | | | Treatment | F (1, 20) = 0.232 | 0.6349 |
| | | | | Genotype × Treatment | F (1, 20) = 0.173 | 0.6821 |
| **1J** | CC-FKBP51 | WT-SAL vs WT-KA | Student's *t* test (unpaired, two-tailed) | Treatment | t (10) = 2.854 | 0.0171 |
| **1K** | HPC-FKBP51 | WT-SAL vs WT-KA | Student's *t* test (unpaired, two-tailed) | Treatment | t (10) = 0.8288 | 0.4265 |
| **1M** | Node density | WT vs *Fkbp5*-KO | Two-way ANOVA with Tukey's *post hoc* test | Genotype | F (1, 21) = 3.12 | 0.0918 |
| | | | | Treatment | F (1, 21) = 63.5 | < 0.0001 |
| | | | | Genotype × Treatment | F (1, 21) = 51.7 | < 0.0001 |
| **1N** | $Na_V1.6$ domain length | WT vs *Fkbp5*-KO | Two-way ANOVA with Tukey's *post hoc* test | Genotype | F (1, 1001) = 11.0 | 0.0010 |
| | | | | Treatment | F (1, 1001) = 24.7 | < 0.0001 |
| | | | | Genotype × Treatment | F (1, 1001) = 144 | < 0.0001 |

**Supplementary Table S4.** Summary of statistical analysis. (continued)

| Figure | Measurement | Group Comparison | Type of test | Factor | Statistical value | *P* value |
|---|---|---|---|---|---|---|
| **2D** | Body weight | *Fkbp5*$^{fl/fl}$ vs. a*Fkbp5*-cKO | Student's *t* test (unpaired, two-tailed) | Genotype | t (10) = 0.5316 | 0.6066 |
| **2E** | Body weight | *Fkbp5*$^{fl/fl}$ vs. a*Fkbp5*-cKO | Two-way ANOVA | Genotype × Time | F (1, 10) = 0.2826 | 0.6066 |
| **2G** | FKBP51 | *Fkbp5*$^{fl/fl}$ vs. a*Fkbp5*-cKO | Student's *t* test (unpaired, two-tailed) | Genotype | t (7) = 4.148 | 0.0043 |
| | GLAST | *Fkbp5*$^{fl/fl}$ vs. a*Fkbp5*-cKO | Student's *t* test (unpaired, two-tailed) | Genotype | t (7) = 1.58 | 0.1582 |
| **2H** | OFT-total distance | WT vs. *Fkbp5*-KO | Student's *t* test (unpaired, two-tailed) | Genotype | t (10) = 0.4719 | 0.6472 |
| | | *Fkbp5*$^{fl/fl}$ vs. a*Fkbp5*-cKO | Student's *t* test (unpaired, two-tailed) | Genotype | t (10) = 0.6877 | 0.5073 |
| **2I** | OFT-total distance | WT vs. *Fkbp5*-KO | Student's *t* test (unpaired, two-tailed) | Genotype | t (10) = 1.348 | 0.2073 |
| | | *Fkbp5*$^{fl/fl}$ vs. a*Fkbp5*-cKO | Student's *t* test (unpaired, two-tailed) | Genotype | t (10) = 0.5582 | 0.5890 |
| **3B** | Seizure score | *Fkbp5*$^{fl/fl}$-KA vs. a*Fkbp5*-cKO-KA | Two-way repeated measures ANOVA followed by Bonferroni's *post hoc* test | Genotype | F (1, 16) = 5.388 | 0.0338 |
| | | | | Treatment | F (12, 192) = 5.810 | < 0.0001 |
| | | | | Genotype × Treatment | F (12, 192) = 2.556 | 0.0037 |
| **3D** | Discrimination index | *Fkbp5*$^{fl/fl}$ vs. a*Fkbp5*-cKO | Two-way repeated measures ANOVA followed by Bonferroni's *post hoc* test | Genotype × Treatment | F (1, 32) = 8.47 | 0.0065 |
| **3F** | GFAP | *Fkbp5*$^{fl/fl}$ vs. a*Fkbp5*-cKO | Two-way ANOVA with Tukey's *post hoc* test | Genotype | F (1, 16) = 11.28 | 0.0040 |
| | | | | Treatment | F (1, 16) = 12.29 | 0.0029 |
| | | | | Genotype × Treatment | F (1, 16) = 20.60 | 0.0003 |
| | GLT-1 | *Fkbp5*$^{fl/fl}$ vs. a*Fkbp5*-cKO | Two-way ANOVA with Tukey's *post hoc* test | Genotype | F (1, 16) = 1.721 | 0.2081 |
| | | | | Treatment | F (1, 16) = 0.2541 | 0.6211 |
| | | | | Genotype × Treatment | F (1, 16) = 0.8222 | 0.3780 |
| **3H** | GFAP area, CA3 | *Fkbp5*$^{fl/fl}$ vs. a*Fkbp5*-cKO | Two-way ANOVA with Tukey's *post hoc* test | Genotype | F (1, 16) = 5.091 | 0.0384 |
| | | | | Treatment | F (1, 16) = 2.393 | 0.1414 |
| | | | | Genotype × Treatment | F (1, 16) = 6.279 | 0.0234 |
| **3J** | GLT-1 area, CA3-SP+SL | *Fkbp5*$^{fl/fl}$ vs. a*Fkbp5*-cKO | Two-way ANOVA with Tukey's *post hoc* test | Genotype | F (1, 16) = 33.7 | < 0.0001 |
| | | | | Treatment | F (1, 16) = 7.25 | 0.0160 |
| | | | | Genotype × Treatment | F (1, 16) = 16.9 | 0.0008 |
| | GLT-1 area, CA3-SR | *Fkbp5*$^{fl/fl}$ vs. a*Fkbp5*-cKO | Two-way ANOVA with Tukey's *post hoc* test | Genotype | F (1, 16) = 43.7 | < 0.0001 |
| | | | | Treatment | F (1, 16) = 3.17 | 0.0939 |
| | | | | Genotype × Treatment | F (1, 16) = 12.2 | 0.0030 |
| **4B** | MAP2 | WT vs *Fkbp5*-KO | Two-way ANOVA with Tukey's *post hoc* test | Genotype | F (1, 12) = 38.9 | < 0.0001 |
| | | | | Treatment | F (1, 12) = 14.8 | 0.0023 |
| | | | | Genotype × Treatment | F (1, 12) = 4.85 | 0.0479 |
| **4C** | GFAP area | WT vs *Fkbp5*-KO | Two-way ANOVA with Tukey's *post hoc* test | Genotype | F (1, 8) = 11.5 | 0.0094 |
| | | | | Treatment | F (1, 8) = 0.0664 | 0.8032 |
| | | | | Genotype × Treatment | F (1, 8) = 57.2 | < 0.0001 |
| **4E** | FKBP51 | WT-CTL vs WT-NMDA | Student's *t* test (unpaired, two-tailed) | Treatment | t (4) = 1.344 | *0.25* |
| **4F** | p-p65/p65 | WT vs *Fkbp5*-KO | Two-way ANOVA with Tukey's *post hoc* test | Genotype | F (1, 8) = 24.2 | 0.0012 |
| | | | | Treatment | F (1, 8) = 14.8 | 0.0049 |
| | | | | Genotype × Treatment | F (1, 8) = 1.76 | 0.2207 |
| **4G** | PHLPP1 | WT vs *Fkbp5*-KO | Two-way ANOVA with Tukey's *post hoc* test | Genotype | F (1, 8) = 183 | < 0.0001 |
| | | | | Treatment | F (1, 8) = 675 | < 0.0001 |
| | | | | Genotype × Treatment | F (1, 8) = 179 | < 0.0001 |
| **4H** | p-AKT/AKT | WT vs *Fkbp5*-KO | Two-way ANOVA with Tukey's *post hoc* test | Genotype | F (1, 8) = 0.725 | 0.4193 |
| | | | | Treatment | F (1, 8) = 12.1 | 0.0083 |
| | | | | Genotype × Treatment | F (1, 8) = 1.67 | 0.2321 |

**Supplementary Table S4.** Summary of statistical analysis. (continued)

| Figure | Measurement | Group Comparison | Type of test | Factor | Statistical value | *P* value |
|---|---|---|---|---|---|---|
| **5A** | *Fkbp5* mRNA | - | One-way ANOVA with Tukey's *post hoc* test | Construct | F (2, 12) = 12.04 | 0.0014 |
| **5C** | Myc-FKBP5 | Myc-*Fkbp5*-WT vs. Myc-*Fkbp5*-3AR | Two-way ANOVA with Tukey's *post hoc* test | Construct | F (1, 20) = 0.714 | 0.4083 |
| | | | | Treatment | F (1, 20) = 0.498 | 0.4887 |
| | | | | Construct × Treatment | F (1, 20) = 0.0277 | 0.8695 |
| **5D** | FKBP51 | Myc vs. Myc-*Fkbp5*-WT | Two-way ANOVA with Tukey's *post hoc* test | Construct | F (1, 16) = 165.7 | < 0.0001 |
| | | | | Treatment | F (1, 16) = 2.755 | 0.1164 |
| | | | | Construct × Treatment | F (1, 16) = 0.3498 | 0.5625 |
| | | Myc vs. Myc-*Fkbp5*-3AR | Two-way ANOVA with Tukey's *post hoc* test | Construct | F (1, 16) = 312.2 | < 0.0001 |
| | | | | Treatment | F (1, 16) = 0.08570 | 0.7735 |
| | | | | Construct × Treatment | F (1, 16) = 1.001 | 0.3320 |
| | | Myc-*Fkbp5*-WT vs. Myc-*Fkbp5*-3AR | Two-way ANOVA with Tukey's *post hoc* test | Construct | F (1, 16) = 2.521 | 0.1319 |
| | | | | Treatment | F (1, 16) = 0.5873 | 0.4546 |
| | | | | Construct × Treatment | F (1, 16) = 1.701 | 0.2106 |
| **5E** | p-p65/p65 | Myc vs. Myc-*Fkbp5*-WT | Two-way ANOVA with Tukey's *post hoc* test | Construct | F (1, 16) = 4.813 | 0.0434 |
| | | | | Treatment | F (1, 16) = 28.28 | <0.0001 |
| | | | | Construct × Treatment | F (1, 16) = 1.407 | 0.2529 |
| | | Myc vs. Myc-*Fkbp5*-3AR | Two-way ANOVA with Tukey's *post hoc* test | Construct | F (1, 16) = 0.4041 | 0.5340 |
| | | | | Treatment | F (1, 16) = 15.31 | 0.0012 |
| | | | | Construct × Treatment | F (1, 16) = 1.098 | 0.3102 |
| | | Myc-*Fkbp5*-WT vs. Myc-*Fkbp5*-3AR | Two-way ANOVA with Tukey's *post hoc* test | Construct | F (1, 16) = 3.707 | 0.0722 |
| | | | | Treatment | F (1, 16) = 26.44 | P<0.0001 |
| | | | | Construct × Treatment | F (1, 16) = 5.673 | P=0.0300 |
| **5F** | *Tnf-α* | Myc vs. Myc-*Fkbp5*-WT | Two-way ANOVA with Tukey's *post hoc* test | Construct | F (1, 16) = 0.7926 | P=0.3865 |
| | | | | Treatment | F (1, 16) = 899.6 | P<0.0001 |
| | | | | Construct × Treatment | F (1, 16) = 1.566 | P=0.2288 |
| | | Myc vs. Myc-*Fkbp5*-3AR | Two-way ANOVA with Tukey's *post hoc* test | Construct | F (1, 16) = 10.79 | P=0.0047 |
| | | | | Treatment | F (1, 16) = 683.9 | P<0.0001 |
| | | | | Construct × Treatment | F (1, 16) = 9.512 | P=0.0071 |
| | | Myc-*Fkbp5*-WT vs. Myc-*Fkbp5*-3AR | Two-way ANOVA with Tukey's *post hoc* test | Construct | F (1, 16) = 12.51 | P=0.0027 |
| | | | | Treatment | F (1, 16) = 535.9 | P<0.0001 |
| | | | | Construct × Treatment | F (1, 16) = 13.52 | P=0.0020 |
| **5G** | *Il-1β* | Myc vs. Myc-*Fkbp5*-WT | Two-way ANOVA with Tukey's *post hoc* test | Construct | F (1, 12) = 2.137 | 0.1695 |
| | | | | Treatment | F (1, 12) = 28.72 | 0.0002 |
| | | | | Construct × Treatment | F (1, 12) = 2.141 | 0.1691 |
| | | Myc vs. Myc-*Fkbp5*-3AR | Two-way ANOVA with Tukey's *post hoc* test | Construct | F (1, 12) = 0.7490 | 0.4038 |
| | | | | Treatment | F (1, 12) = 39.30 | < 0.0001 |
| | | | | Construct × Treatment | F (1, 12) = 0.7522 | 0.4028 |
| | | Myc-*Fkbp5*-WT vs. Myc-*Fkbp5*-3AR | Two-way ANOVA with Tukey's *post hoc* test | Construct | F (1, 12) = 0.8075 | 0.3865 |
| | | | | Treatment | F (1, 12) = 41.21 | 0.0001 |
| | | | | Construct × Treatment | F (1, 12) = 0.8075 | 0.3865 |
| **5I** | p-AKT/AKT | Myc vs. Myc-*Fkbp5*-WT | Two-way ANOVA with Tukey's *post hoc* test | Construct | F (1, 12) = 1.503 | 0.2437 |
| | | | | Treatment | F (1, 12) = 2.075 | 0.1753 |
| | | | | Construct × Treatment | F (1, 12) = 1.655 | 0.2225 |
| | | Myc vs. Myc-*Fkbp5*-3AR | Two-way ANOVA with Tukey's *post hoc* test | Construct | F (1, 8) = 0.009219 | 0.9259 |
| | | | | Treatment | F (1, 8) = 1.058 | 0.3337 |
| | | | | Construct ×Treatment | F (1, 8) = 4.757 | 0.0608 |
| | | Myc-*Fkbp5*-WT vs. Myc-*Fkbp5*-3AR | Two-way ANOVA with Tukey's *post hoc* test | Construct | F (1, 12) = 1.398 | 0.2600 |
| | | | | Treatment | F (1, 12) = 0.1391 | 0.7157 |
| | | | | Construct × Treatment | F (1, 12) = 0.2622 | 0.6179 |

**Supplementary Fig. S1 Uncropped images of Western blots related to figure 1 and 2.**

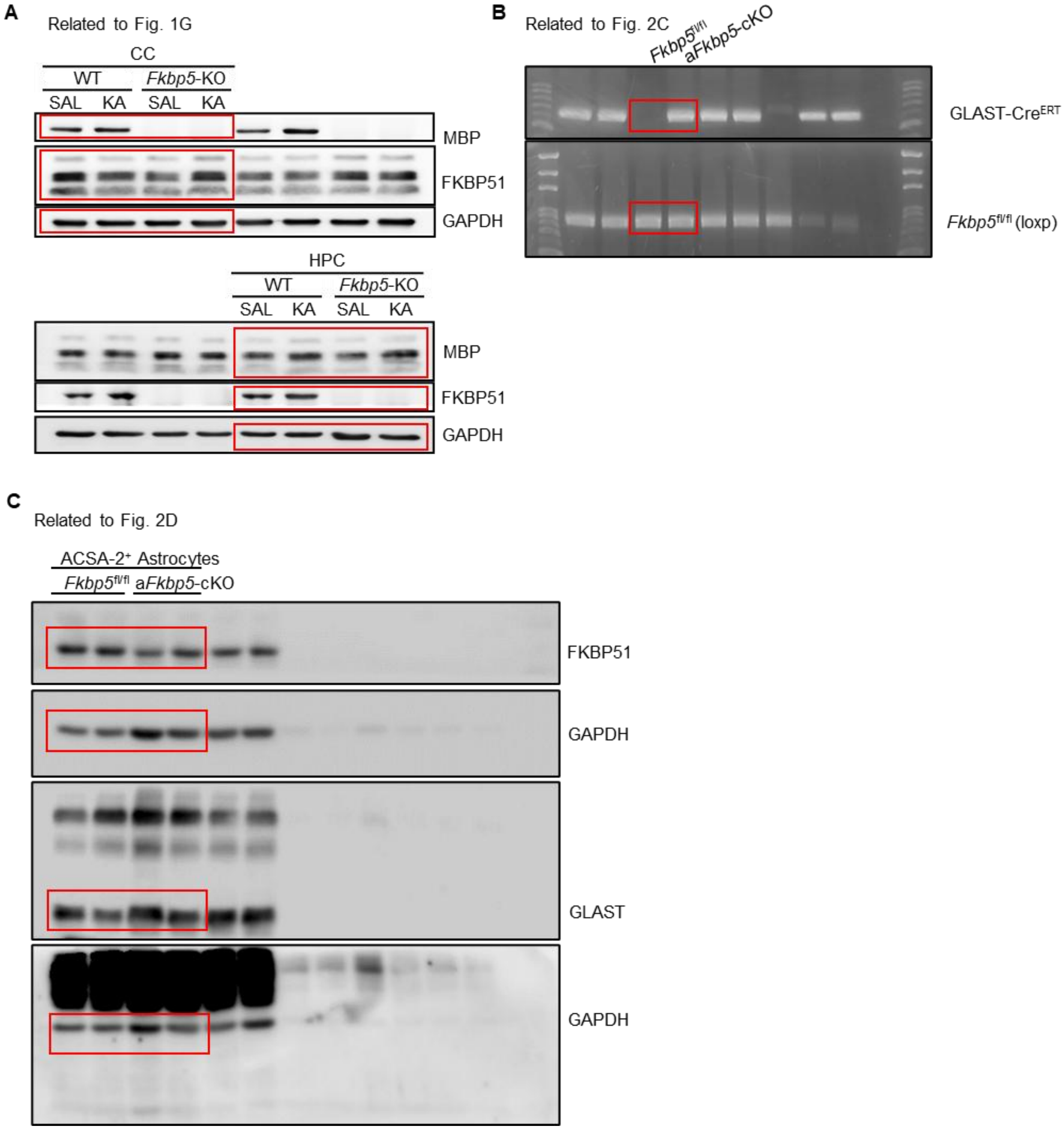

**Supplementary Fig. S2 Uncropped images of Western blots related to figure 3, 4, and 5.**

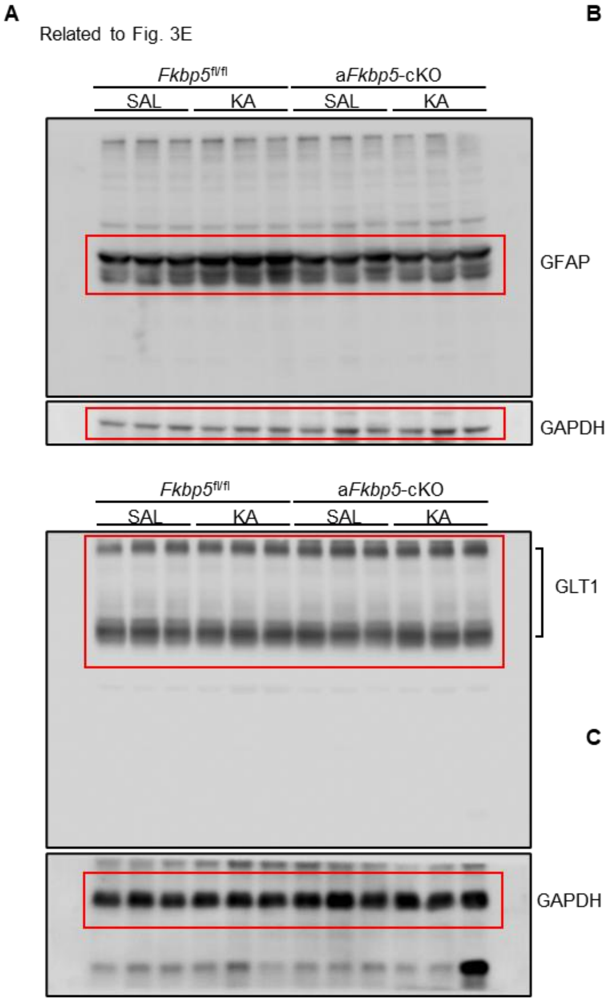


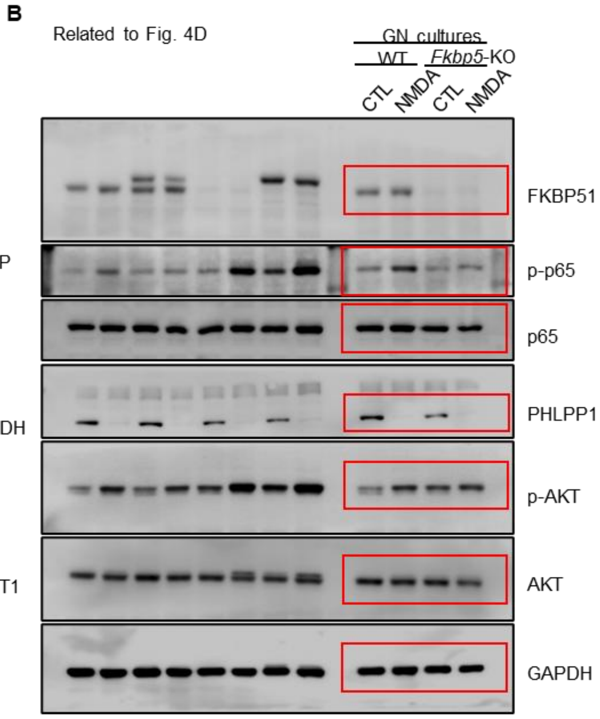


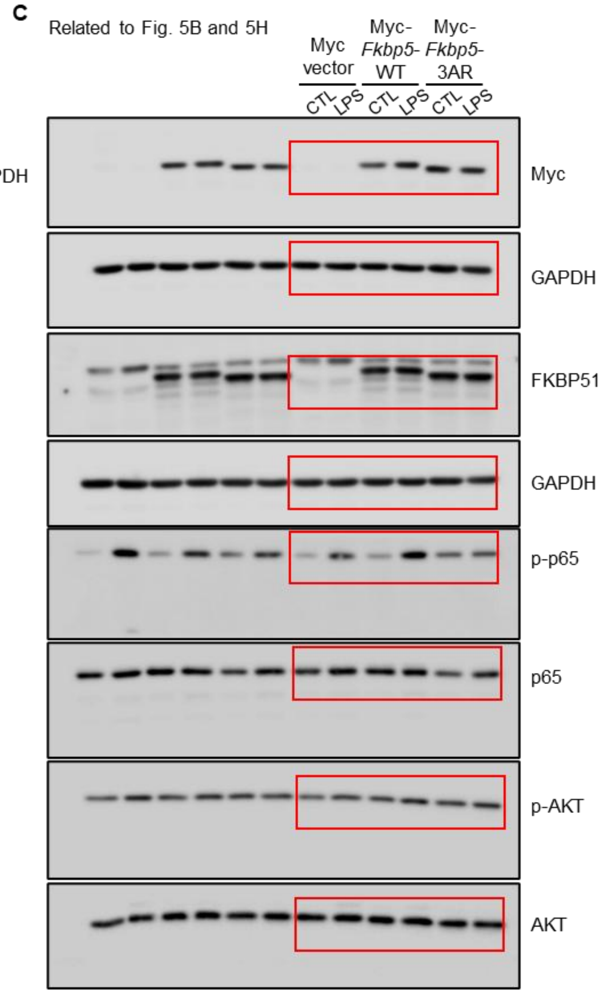

**Supplementary Fig. S3 *Fkbp5* deletion reverses AMPA-induced damage to oligodendrocytes.**

Primary WT and *Fkbp5*-KO oligodendrocytes (OLs) were treated with AMPA for 24 hours. **A** Representative immunofluorescence images of Rip (red)- and TUNEL (green)-stained cells counterstained with DAPI (blue). Scale bar: 30 μm. **B-D** Quantitative results for cell death (B), $Rip^+$ process length (C), and average process intersections (D). *n* = 5 **E, F** RT–qPCR results for *Fkbp5* (E) and *Mbp* (F) mRNA expression in AMPA-treated OLs. *n* = 6. The data are expressed as the mean ± SEM. **P* < 0.05, ***P* < 0.01, and ****P* < 0.001 compared between the two indicated groups by two-way ANOVA with Tukey's *post hoc* test.

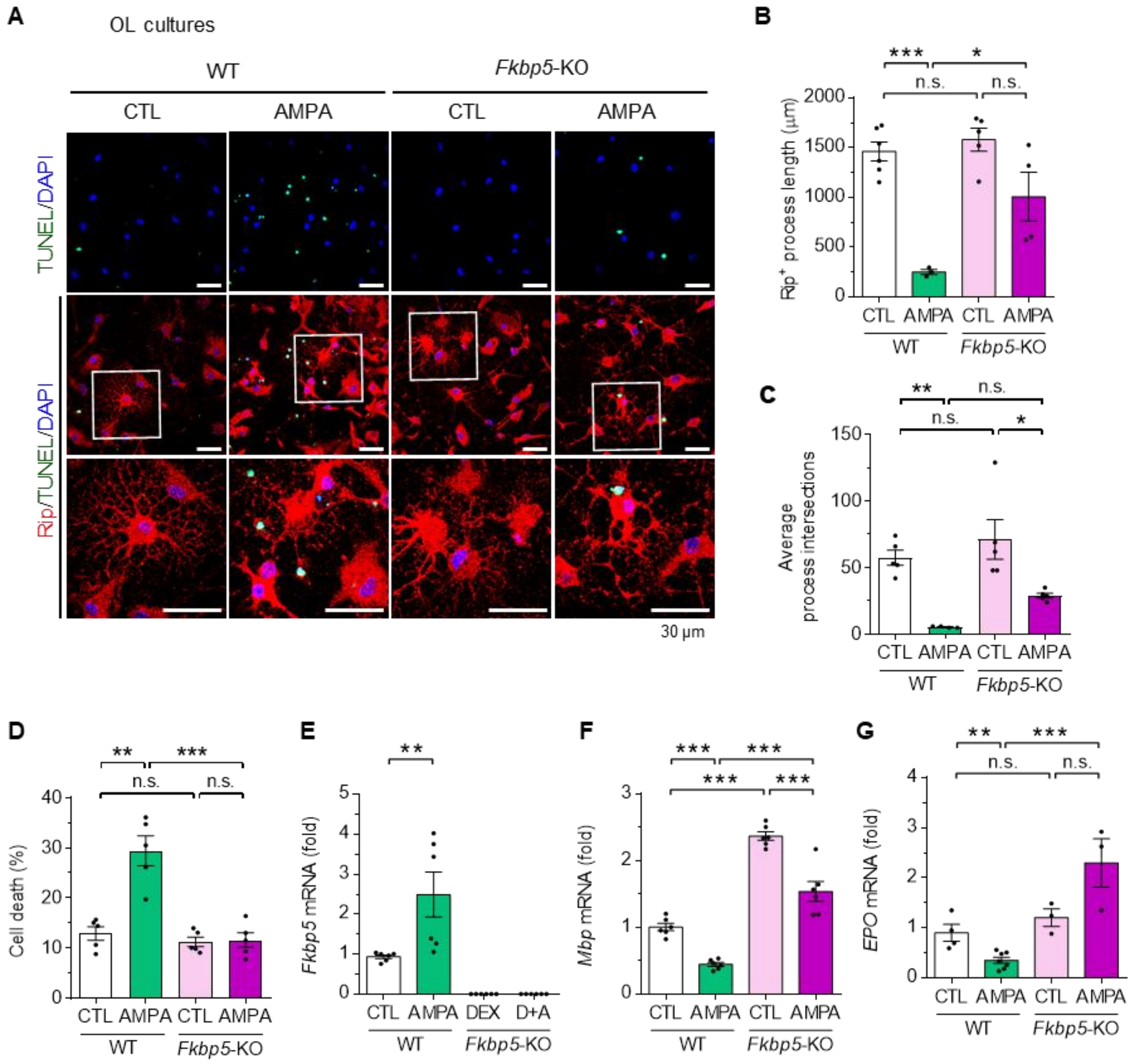

### Materials and Methods for Supplementary Fig. S3

#### *Primary culture of mouse oligodendrocytes*

Primary oligodendrocytes (OLs) were prepared from postnatal day 1 to 2 mice as previously described [1]. The cultured cell purity was > 90% on the basis of receptor-interacting protein (RIP) immunoreactivity.

#### *TUNEL staining for cell apoptosis*

Cell apoptosis was analyzed using the TUNEL assay with an In Situ Cell Death Detection Kit (Roche). The cells were washed with PBS, fixed in 4% formaldehyde (20 minutes, RT), and permeabilized with 0.1% Triton X-100/0.1% sodium citrate (2 minutes, 4°C). TUNEL staining was performed following the manufacturer's protocol. Images were captured using an Olympus FV1000 laser confocal microscope. Apoptotic cells were identified as TUNEL-positive with condensed DAPI-stained nuclei, and total DAPI-stained cells served as the reference. Three randomly selected areas per well were counted, with three independent wells per condition. Cell death was calculated as the percentage of TUNEL-positive cells relative to the total cell count.

#### *Reverse transcription–quantitative polymerase chain reaction (RT–qPCR)*

Total mRNA was extracted using TRIzol reagent (Invitrogen) in accordance with the manufacturers' instructions. Total mRNA was reverse transcribed into cDNA using a high-capacity reverse transcription kit (Thermo Fisher Scientific). Quantitative PCR was performed using an ABI StepOnePlus real-time PCR system (Applied Biosystems) with 2x SYBR Green PCR Master Mix (Applied Biosystems). The sequences of primers used for RT–qPCR are listed in Supplementary Table 3. The difference between the threshold cycle (Ct) values for the target and reference genes in each sample was calculated. The relative fold change in gene expression was determined using the ΔΔCt method and expressed as $2^{-\Delta\Delta Ct}$.
